# Balancing Activation and Repression: CoREST-p300 Antagonism Controls Retinoic Acid-Driven Differentiation in AML

**DOI:** 10.1101/2025.10.13.682235

**Authors:** Mina M. Tayari, Helena Gomes Dos Santos, Sadat Dokaneheifard, Samuel D. Whedon, Monica Valencia, Harikumar Arigela, Felipe Beckedorff, Justin M. Watts, Philip A. Cole, Ramin Shiekhattar

## Abstract

The histone demethylase KDM1A (LSD1), a component of the CoREST corepressor complex, is highly expressed in hematologic malignancies and regulates hematopoietic differentiation. Despite its essential developmental role, LSD1 inhibition has emerged as a promising strategy to enhance retinoic acid (RA)-responsive gene expression in subsets of acute myeloid leukemia (AML). Here, we show that LSD1 physically interacts with RAR/RXR heterodimers at specific genomic loci, restricting chromatin accessibility and transcriptional activation of differentiation programs. Single-agent inhibition of LSD1 or HDACs promotes only partial differentiation. In contrast, Corin, a dual LSD1/CoREST inhibitor, synergizes with all-trans retinoic acid (ATRA) to induce robust myeloid differentiation and apoptosis. Corin treatment alone does not significantly increase H3K4me3 levels; however, in combination with ATRA, it disrupts CoREST-RAR/RXR complexes and facilitates the recruitment of the coactivator p300. Together, they shift chromatin to an active state, enhancing H3K4me3 via increased transcriptional engagement and coactivator recruitment. Our findings identify the functional antagonism between CoREST and p300 as a regulatory axis of RA signaling in AML. Targeting this mechanism with Corin and ATRA re-sensitizes non-APL AML cells to RA-induced differentiation, suggesting a broader therapeutic approach for overcoming resistance in ATRA-refractory leukemias.

**Significance:** Co-inhibition of LSD1 and HDACs with Corin, combined with ATRA, reprograms chromatin by replacing CoREST with p300, uniquely activating differentiation-associated genes and offering a targeted strategy for treating AML. *Graphical abstract created with BioRender* (https://biorender.com).

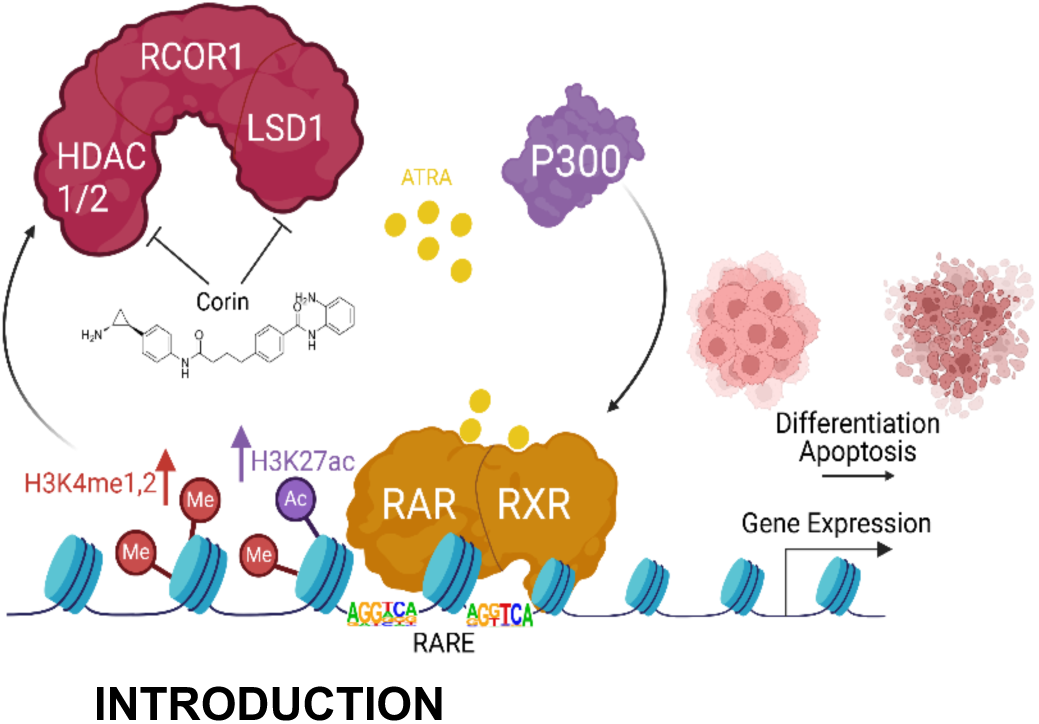

## INTRODUCTION

Treatment of acute myeloid leukemia (AML) typically involves intensive chemotherapy regimens designed to eradicate leukemia cells in the blood and bone marrow. Despite these efforts, considerable risk remains that the disease may relapse or develop resistance following the treatments. We and others previously showed that all-trans retinoic acid (ATRA) in combination with epigenetic drugs such as Lysine specific demethylase 1 (LSD1) inhibitors unlock the ATRA-driven therapeutic response pinpointing to LSD1 as a new therapeutic target [1–4]. LSD1 is a histone demethylase implicated in the maintenance of pluripotency and proliferation and LSD1 expression is known to be upregulated in myeloid malignancies [5]. LSD1 is a key component of the corepressor for element 1-silencing transcription factor (CoREST) complex, which plays a crucial role in chromatin remodeling and gene regulation. The CoREST complex, also including components like RCOR1, 2, or 3 (also known as CoREST) and histone deacetylase HDAC 1/2, which interact with LSD1 to form a functional unit that can modify chromatin structure and regulate transcriptional activity. The acetylation and methylation of lysine residues in the tails of histone proteins are critical not only for recruitment of co-activators and co-repressors but also for shaping the structure of chromatin itself. Beyond demethylase activity, the LSD1 component of the CoREST complex has been shown to be recruited to chromatin by SNAG family transcription factors (e.g., SNAIL, GFI1) in a manner that is competed by most LSD1 active site inhibitor [6]. The Retinoic Acid Receptor (RAR) functions as a nuclear receptor and ligand-activated transcription factor. RAR typically forms heterodimers with RXR and binds to retinoic acid response elements (RAREs) in DNA [7]. RAR-RXR regulates transcription by binding to RARE found in the enhancer regions of RA-responsive genes [8]. Retinoic acid (RA) signaling via the RAR/RXR pathway triggers differentiation, cell cycle arrest, and apoptosis in various cell types.

In the classical model, the presence of RA as a ligand facilitates the recruitment of coactivators and the release of corepressors, thereby enhancing the transcription of target genes [9–11]. Conversely, in the absence of RA, RAR/RXR binds to corepressor proteins NcoR1[12] and SMRT [13], resulting in the suppression of transcription. LSD1 is involved in both corepressor and coactivator complexes, contributing to the regulation of certain transcription factors including nuclear receptors, such as estrogen receptor α (ER α) [14, 15] and androgen receptor (AR) [16]. However, the role of LSD1 in the dysregulation of RAR/RXR signaling remains unclear. LSD1 might affect the chromatin structure at the RAREs and modulate the accessibility of the target genes to transcriptional machinery. Despite these insights, further research is necessary to fully elucidate how LSD1 contributes to RAR/RXR signaling and target gene activation in leukemia.

Previously, we and others showed that RA signaling can be enhanced by combining ATRA with LSD1 inhibitors [1–3]. There were also reports of other epigenetic therapies used in combination with HDAC inhibitors [17, 18]. Recently, Corin was identified as a potent and selective dual inhibitor that targets both LSD1 and HDAC1/2, which are key components of the CoREST complex [19]. Additionally, Corin has demonstrated effectiveness in inhibiting the growth of various cancers, including melanoma [19, 20], cutaneous squamous cell carcinoma [19], diffuse intrinsic pontine glioma [21] and breast cancers [22]. Here, we utilized Corin in combination with ATRA to induce differentiation and epigenetic reprogramming in leukemia. These findings highlight the interplay between epigenetic modifiers and RAR signaling, revealing their transcriptional impact on RA-responsive genes and the resulting differentiation of leukemic cells. Our results highlight a mechanism by which CoREST inhibition facilitates the recruitment of P300 to RAR target genes, thereby enhancing transcriptional activation and promoting differentiation.

## RESULTS

### Co-targeting HDACs and LSD1 with Corin Synergistically reduce AML Growth *In Vitro*

To better understand the therapeutic potential of dual targeting of HDACs and LSD1, we assessed the effects of Corin in AML. We treated MOLM-13 cells with the dual inhibitor Corin (0.1 µM). Additionally, we used Iadademstat (ORY-1001, 0.1 µM), a potent LSD1 inhibitor; or Entinostat (0.1 µM), a class I-selective HDAC inhibitor alone or in combination. We performed RNA sequencing (RNA-seq) to analyze transcriptomic changes after treating MOLM-13 cells for 24 hours with DMSO (control), ORY-1001, Entinostat, the combination of ORY-1001 and Entinostat, or Corin alone **(Fig. 1A)**. Treatment with ORY-1001 alone resulted in only minor gene expression changes compared to the control **(Fig. 3A)**, consistent with prior observations that MOLM-13, an acute monocytic leukemia (AML) cell line, is resistant to LSD1 inhibition [23]. In contrast, treatment with Entinostat alone resulted in substantial gene expression changes. However, the most pronounced transcriptional alterations occurred with the combination of ORY-1001 and Entinostat, amplifying both upregulated and downregulated gene expression profiles.

**Figure 1.**
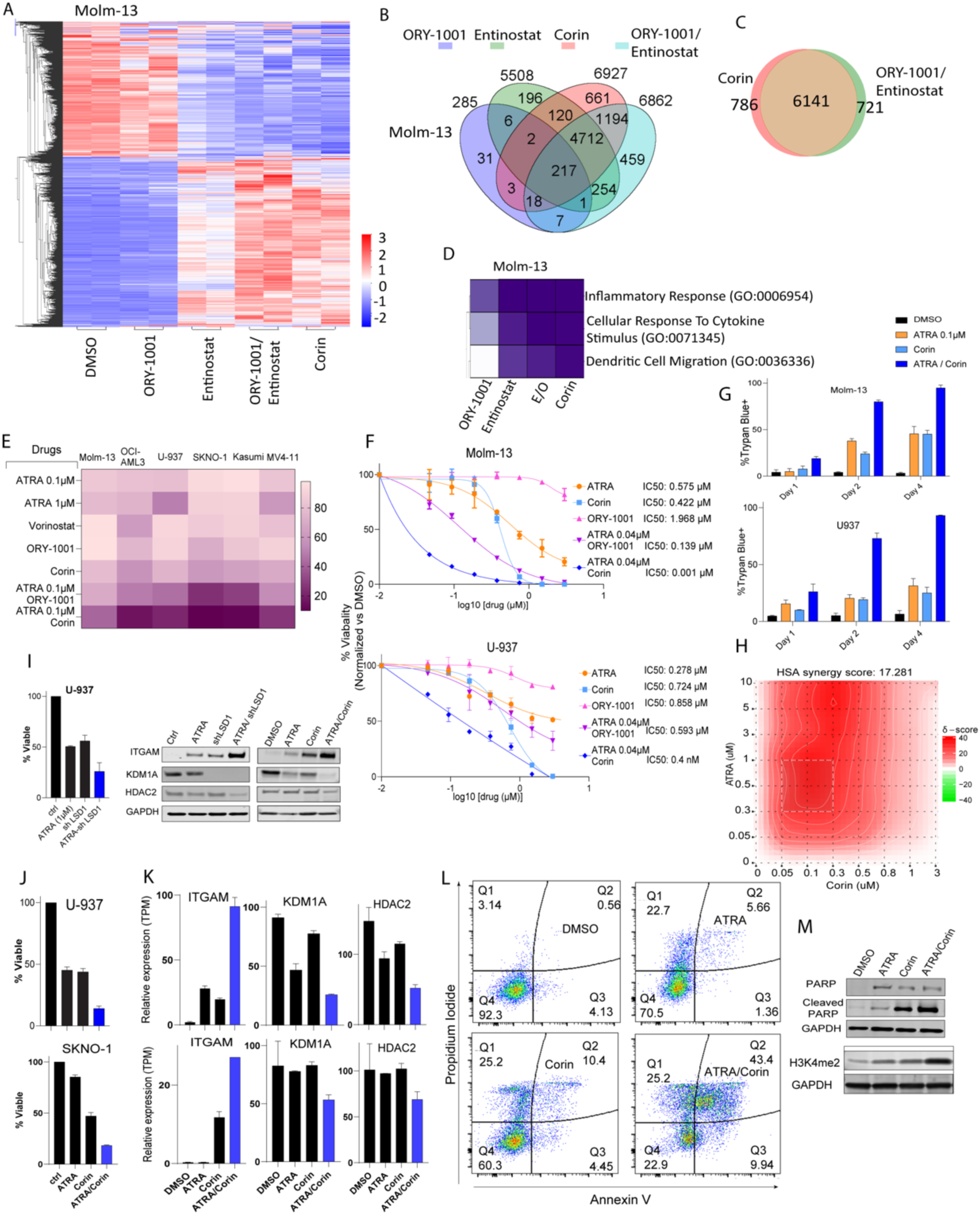
Corin phenocopies combined LSD1 and HDAC inhibition in gene expression and shows the strongest inhibitory effect on AML cell proliferation compared to LSD1 or HDAC inhibition alone. **A)** Heatmap illustrating gene expression in MOLM-13 cells treated for 24 hours with Corin, compared to treatments with the LSD1 inhibitor ORY-1001 (0.1 µM) or the HDAC inhibitor Entinostat (0.1 µM) (alone or in combination). **B)** Venn diagram displaying the overlap of differentially expressed genes between Corin, ORY-1001, and Entinostat treatments (single-agent treatments or combination) in MOLM-13 cells. **C)** Ven diagram displaying the overlap of differentially expressed genes between Corin and ORY-1001/Entinostat treatment in MOLM-13. **D)** Heatmap depicting gene ontology pathways activated by treatment with Corin, ORY-1001 and/or Entinostat, compared to DMSO control. **E)** Percent of cell viability relative to DMSO control. **F)** IC50 values of MOLM-13 and U-937 cells treated with low-dose ATRA, ORY-1001 (LSD1 inhibitor), Corin, and their combinations with low-dose ATRA. **G)** Percentage of Trypan blue-positive cells indicating cell death at 2 and 4 days post-treatment. **H)** Synergy score of ATRA-Corin combination treatment. **I)** Cell viability upon LSD1 knockdown using siRNA and its combination with ATRA in U-937 cells. **J)** Cell viability of U-937 cells treated with ATRA-Corin combination. **K)** mRNA expression levels of *ITGAM* (CD11b), *KDM1A*, and *HDAC2* in U-937 cells. **L)** Apoptosis analysis by Annexin V/7AAD staining in U-937 cells. **M)** Western blot analysis of PARP and cleaved PARP following treatment with ATRA, Corin, and their combination, along with changes in H3K4me2 levels.

Notably, Corin alone produced gene expression changes highly similar to those of combination therapy, supporting the conclusion that Corin effectively replicates the combined effects of ORY-1001 and Entinostat **(Fig. 1A)**. While treatment with ORY-1001 alone resulted in significant gene expression changes in only 285 genes, Entinostat alone impacted a larger set of 5,508 genes **(Fig. 1B)**. Corin treatment triggered a broad transcriptional effect, significantly altering 6,927 genes (FDR < 0.05, 2-fold; **Fig. 1B**). Interestingly, we observed a substantial overlap of 6,141 genes between cells treated with the combination of ORY-1001 plus Entinostat and those treated with Corin alone **(Fig. 1C)**. Gene ontology (GO) analysis shows similar pathway activation by Corin and the combination of ORY-1001 and/or Entinostat **(Fig. 1D)**. Genes upregulated by Corin were notably involved in cellular differentiation and inflammatory response pathways **(Fig. S1A-B)**. In MOLM-13 and OCI-AML3 cells, Corin and ORY-1001/Entinostat treatments yielded 2,950 and 3,227 differentially expressed genes, respectively. Notably, differentiation-associated genes were significantly upregulated by both treatments in both cell lines **(Fig. S1C-D)**.

We and others have previously demonstrated that combining LSD1 inhibition with ATRA sensitizes AML cells towards differentiation [1, 2, 23]. We next explored whether simultaneously targeting HDACs and LSD1, along with ATRA treatment, could further enhance differentiation in AML. To test this, we assessed drug sensitivity across six genetically diverse AML cell lines harboring MLL rearrangements (MOLM-13, MV4-11), the AML1-ETO translocation (SKNO-1, KASUMI-1), NPM1/DNMT3A mutations (OCI-AML3), and a TP53 mutation (U-937). Cells were treated with either a low dose (0.04 µM) or high dose (1 µM) of ATRA, ORY-1001 (0.1 µM), Entinostat (0.1 µM), and Corin (0.1 µM). Treatments with the LSD1 inhibitors ORY-1001 or Corin were applied either alone or in combination with a low-dose of ATRA **(Fig. 1E)**. At 0.1 µM, Corin reduced viability in all AML cell lines more effectively than ORY-1001 or Entinostat alone, and its combination with low-dose ATRA further enhanced growth inhibition in all AML cells tested **(Fig. 1E)**. MOLM-13 cells are resistant to LSD1 inhibition [23]. Accordingly, ORY-1001 alone had minimal impact on viability, exhibiting a high IC₅₀ and only modest reduction in cell survival **(Fig. 1F)**. Consistent with previous reports and our own data, low-dose ATRA sensitized MOLM-13 cells to LSD1 inhibition, lowering the ORY-1001 IC₅₀ to 139 nM. Corin alone showed an IC₅₀ of 400 nM, but in combination with low-dose ATRA, its IC₅₀ dropped dramatically to 1 nM. A similar effect was observed in U-937 cells: the ATRA-Corin combination reduced viability and increased Trypan blue-positive cells at days 2 and 4 post-treatment **(Fig. 1G)**, with an HSA synergy score of 17.28 **(Fig. 1H)**.

Importantly, shRNA-mediated LSD1 knockdown reduced U-937 cell viability to a level comparable with Corin treatment and enhanced sensitivity to ATRA, as shown by decreased proliferation and upregulation of the differentiation marker CD11b **(Fig. 1I-K)**. ATRA and Corin each induced a modest reduction in LSD1 protein levels; however, their combination resulted in a near-complete depletion of LSD1, as demonstrated by Western blot analysis **(Fig. 1I)**. Consistent with these findings, RNA-seq analysis showed upregulation of *CD11B*, accompanied by reduced expression of *LSD1* and *HDAC2* **(Fig. 1K)**. To assess whether Corin induces apoptosis in U-937 cells, we performed flow cytometry analysis using Annexin V/7-AAD staining. Treatment with ATRA, Corin, or the ATRA-Corin combination resulted in a progressive increase in late apoptotic cells, from 0.56% in control cells to 5.66% with ATRA, 10.4% with Corin, and 43.4% with the combination **(Fig. 1L)**. These results indicate that the ATRA-Corin combination promotes both differentiation and apoptosis in U-937 cells, as evidenced by elevated levels of cleaved PARP, a marker of apoptosis, particularly in the combination-treated group **(Fig. 1M)**. ATRA-Corin treatment increased global H3K4me2 levels **(Fig. 1M)** and CUT&RUN analysis revealed enhanced H3K4me2 enrichment at promoters of differentiation-associated genes (**Fig. S1E)**.

### ATRA Combined with LSD1/HDAC Inhibition Promotes Myeloid Gene Activation Through Chromatin Accessibility and H3K4me2 Broadening

U-937 and SKNO-1 cells were treated for 24 hours with DMSO, Corin alone, or the ATRA-Corin combination. Combination treatment markedly increased the number of differentially expressed genes **(Fig. 2A, Fig. S2A)**. RNA-seq analysis in U-937, revealed a distinct gene expression signature uniquely induced by the combination, with 1,691 genes exclusively upregulated and largely unresponsive to either agent alone **(Fig. 2B)**. Notably, this signature included genes implicated in hematopoietic lineage commitment and differentiation [1, 24]. Gene ontology analysis of these combination-specific targets revealed the top 10 enriched biological processes, including NF-κB signaling pathways **(Fig. 2C)**. This gene set also included mRNAs of differentiation markers such as ITGAM (CD11b) and CD86 **(Fig. 2D)**. The resulting gene expression profile overlapped with a previously reported ATRA-LSD1 sensitivity signature [2], including downregulation of oncogenic drivers such as *MYC* and *MTOR* **(Fig. S2B-D)**.

**Figure 2:**
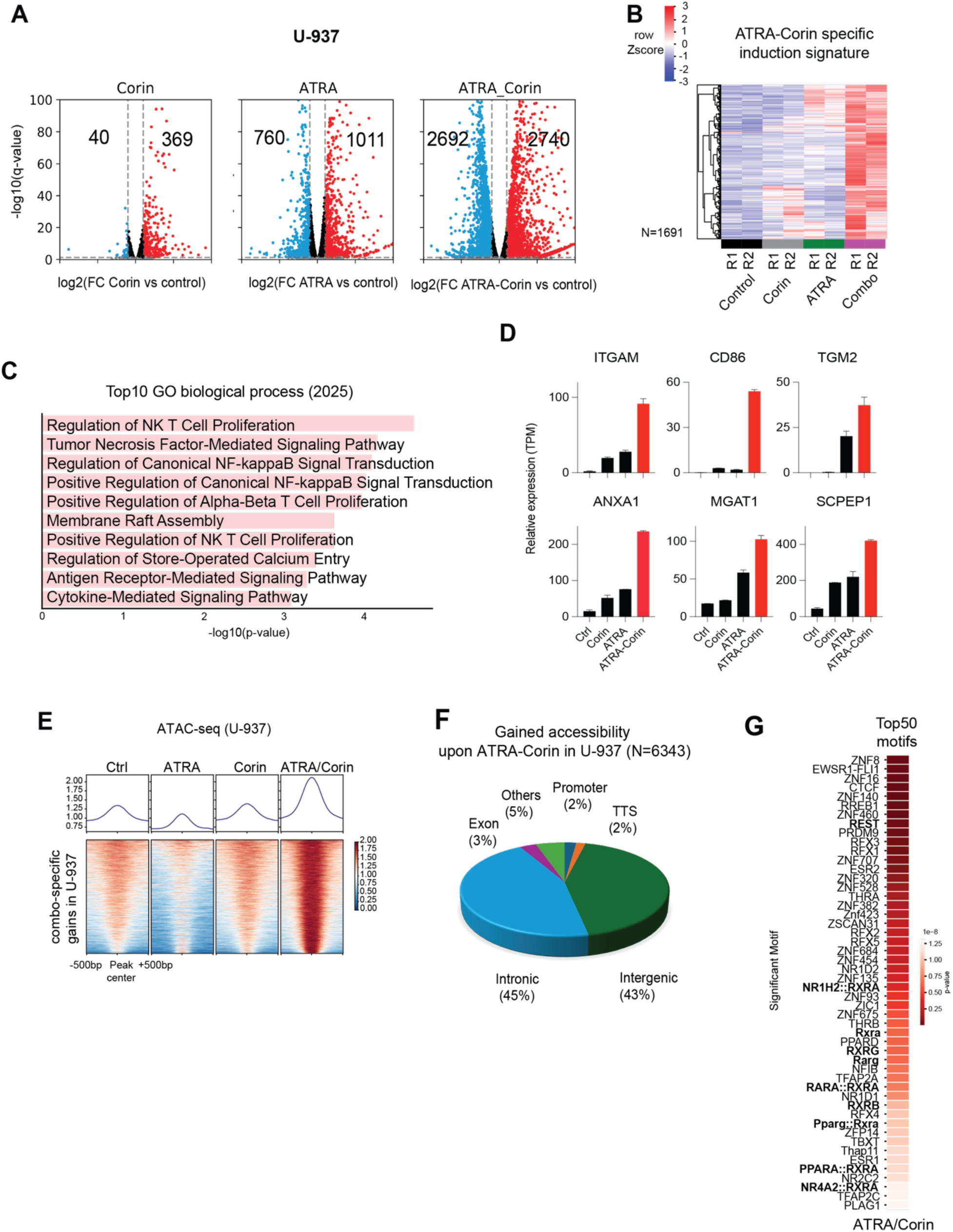
Extensive ATRA/Corin-specific gene induction and gained chromatin accessibility at putative enhancers. **A)** Volcano plots showing differential expressed genes in U-937 upon treatment compared with the untreated DMSO control cells (treatments from left to right: Corin, ATRA, ATRA-Corin; 2-fold change cutoff, and FDR < 0.05, blue = downregulated genes, red = upregulated genes). **B)** Heatmap of the Z-score normalized gene expression in U-937 for a subset of genes obtained by unsupervised clustering of the upregulated genes with positive enrichment in the ATRA-Corin induction signature. **C)** Gene ontology analysis of the top ten biological process in U-937 cells following ATRA-Corin treatment. **D)** Examples of ATRA-Corin responsive genes. **E)** ATRA-Corin Combo specific gain peaks by ATAC-seq **F)** Gained accessibility genome wide distribution upon ATRA-Corin treatment. **G)** Top 50 Significant motifs with gained accessibility.

ATAC-seq analysis showed widespread increases in chromatin accessibility following combination treatment, at putative enhancer regions located within intronic (45%) and intergenic (43%) regions **(Fig. 2E, 2F)**. Notably, motif enrichment analysis identified RAR motifs among the top activated elements **(Fig. 2G)**, which were insufficiently activated by ATRA alone. In contrast, the ATRA-Corin combination selectively enhanced accessibility at RAR-RXR motifs, suggesting a cooperative mechanism involving epigenetic remodeling and retinoid signaling.

We next examined whether the observed transcriptional activation was associated with changes in H3K4me2 distribution (**Fig. S3A-B**). Rather than a uniform increase in signal intensity, ATRA-Corin treatment induced a striking broadening of H3K4me2 peaks across the genome. This widespread expansion of H3K4me2 domains (**Fig. S4A-B**) was accompanied by elevated expression of representative genes such as *TGM2*, *CD58*, and *KLF6* (**Fig. S4C-E**). Notably, regions harboring retinoic acid response elements (RAREs; shown as red blocks) exhibited particularly pronounced H3K4me2 broadening, including at the *RARB* promoter (**Fig. S4F-G**).

### LSD1 and RARα interact physically and cooperatively regulate gene sets critical to AML pathogenesis

We established HEK293 cells stably expressing Flag-tagged retinoic acid receptors (RARs), RARα and RARβ. Using these cell lines, we performed immunoprecipitation (IP) assays and analyzed protein interactions separately in nuclear and cytoplasmic fractions. Mass spectrometry identified known RAR-interacting partners, including NCOA2, NCOA3, and RXRβ. Notably, nuclear fractions also revealed that KDM1A (LSD1), HDAC2, core components of the CoREST complex, as well as the transcriptional coactivator P300, co-immunoprecipitated and associate with both RARα and RARβ (**Fig. 3A-B** and **Table S2)**. Western blotting and silver staining in HEK293T cells confirmed successful purification of RARα and RARβ and validated their interaction with these binding partners **(Fig. 3A-C)**. These findings highlight a dynamic regulatory mechanism involving chromatin coactivators and corepressors, histone modification, and transcriptional activation.

**Figure 3.**
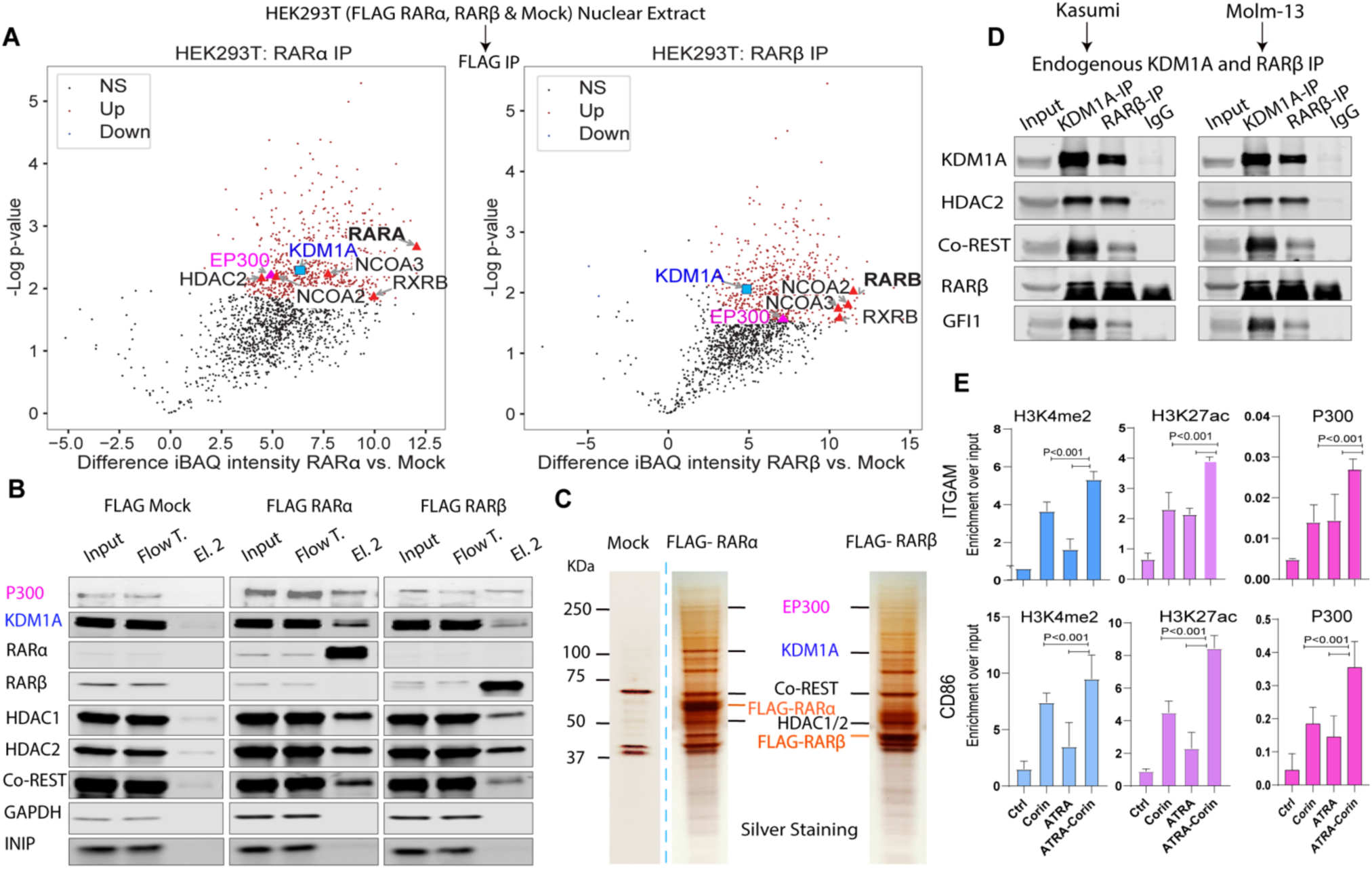
RARα and RARβ interact with LSD1 or the coactivator p300. **A)** Volcano plot illustrating proteins enriched in nuclear extracts from HEK293T cells stably expressing Flag-tagged RARα or RARβ versus mock control (proteins are color-coded as non-significant in black, enriched in red and depleted in blue). Known interacting proteins of RARα and RARβ are indicated with black labels, while LSD1 and p300 are highlighted. **B)** Western blot validation and **C)** silver staining of affinity-purified Flag-RARα and Flag-RARβ complexes from nuclear lysates of stable HEK293T cells. Mock cells served as controls. **D)** Endogenous co-immunoprecipitation confirming interactions between RARβ and LSD1 in KASUMI-1 and MOLM-13 AML cells. **E)** ChIP-qPCR analysis demonstrating increased recruitment of the coactivator p300 at *ITGAM* promoter following ATRA-Corin treatment, accompanied by enrichment of H3K4me2 and H3K27ac at the promoter of ITGAM (top) and CD86 (bottom). Error bars indicate SD from triplicate experiments, and *P* values were determined by a two-tailed unpaired Student’s *t* test.

To validate these interactions in a more physiological context, we performed reciprocal co-immunoprecipitation (co-IP) experiments targeting endogenous RARβ and LSD1 proteins in AML cell lines. These reciprocal co-IP assays confirmed the native interaction between RARβ and LSD1 in MOLM-13 and KASUMI-1 cells **(Fig. 3D)**. Expanding upon these results, we conducted chromatin immunoprecipitation followed by quantitative PCR (ChIP-qPCR) assays to evaluate chromatin modifications at differentiation-related genes upon treatment with Corin and ATRA **(Fig. 3E)**. Combined treatment with ATRA and Corin led to significantly increased enrichment of P300 at *ITGAM*, as well as at the *CD86* promoter **(Fig. 3E)**. This suggests that p300 plays an important role in ATRA-mediated activation of gene expression.

ATRA-Corin treatment led to a marked increase in activating histone marks, including H3K4me3 and H3K27ac, at these regions **(Fig. S5A-B)**. While Corin alone increased H3K27ac at the promoters of these genes, the induction of H3K4me3 and subsequent gene activation required ATRA, underscoring the cooperative effect of the combination treatment. As indicated by red arrows , enhancer regions upstream of *ITGAM* and an intergenic enhancer near *CD86* show LSD1 occupancy accompanied by an increase in H3K4me3 (**Fig. S5A-B**). Similar patterns are observed at the enhancer of *CTSB* and at the promoters of differentiation-associated genes *LY96*, *LILRA5*, and *CD1D* (**Fig. S5C-F**). Collectively, these findings suggest that ATRA-Corin treatment promotes the displacement of LSD1 and the CoREST repressor complex by the coactivator P300 at RAR-responsive loci, facilitating enhanced H3K27 acetylation and robust transcriptional activation of genes essential for AML cell differentiation.

### Global alterations in histone marks in AML Cells treated with Corin or ATRA-Corin

To evaluate changes in histone modifications following treatment, we first confirmed by western blot a global increase in H3K4me2 upon LSD1 inhibition with Corin, which was further enhanced when combined with ATRA **(Fig. 1I)**. CUT&RUN analysis showed that Corin treatment significantly increased the number of peaks detected for H3K4me2 and H3K27ac, whereas minimal changes were observed for H3K4me3 **(Fig. 4A)**. Genome-wide analysis revealed nearly 20,000 Corin-specific gained peaks predominantly reflecting modifications associated with the CoREST complex, primarily H3K4me2 and H3K27ac, with minimal impact on H3K4me3 **(Fig. 4B)**. Further analysis of genomic distribution showed these Corin-specific peaks were mostly located within intronic regions (47.5%), promoters (22.2%), intergenic regions (19.7%), transcription termination sites (TTS; 5.8%), and exons (4.8%) **(Fig. 4C)**. Further analysis showed that LSD1 is located at Corin-specific peak regions **(Fig. 4D)**. Interestingly, while H3K4me2 and H3K27ac marks increased with Corin alone, the elevation of H3K4me3 required additional stimulation by ATRA **(Fig. 4E)**. This aligns with our findings suggesting that LSD1 inhibition disrupts the CoREST complex, resulting in elevated H3K4me2 and H3K27ac, whereas accumulation of H3K4me3 is dependent on a co-activator (e.g., P300) recruitment and activation facilitated by ATRA.

**Figure 4.**
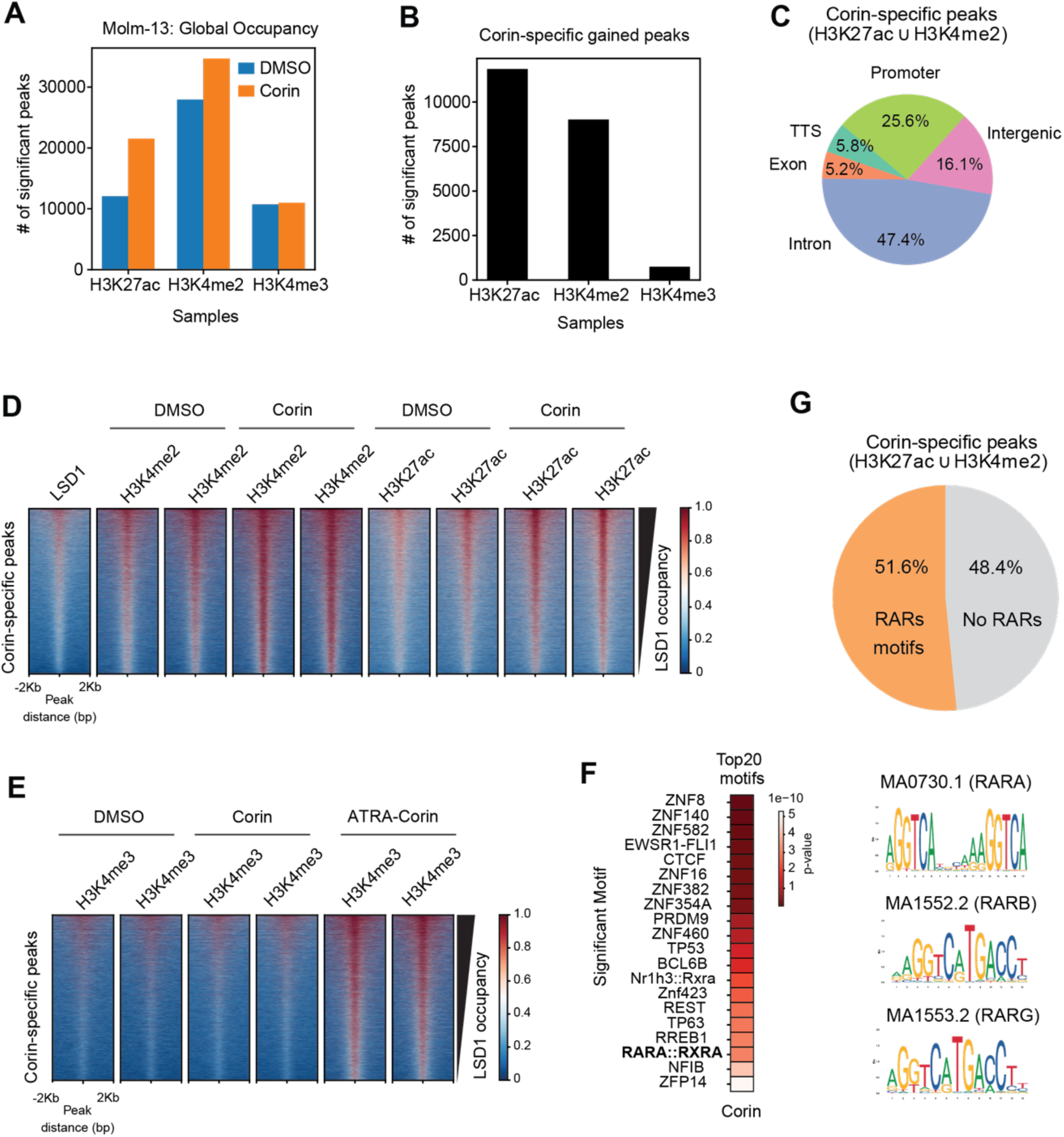
Corin-Induced epigenetic changes mark RAR-enriched regions primed for activation by ATRA. **A)** CUT&RUN peak distribution for the epigenetic marks H3K4me3, H3K27ac, and H3K4me2 in MOLM-13 cells treated with Corin compared to DMSO control for 24 hours. **B)** Bar plot distribution of significant Corin-specific peaks gained after treatment. **C)** Pie chart showing the distribution of Corin-activated peaks across different genomic regions (N=19466 peaks obtained by the union of the common gains across 2 replicates each from H3K4me2 and from H3K27ac CUT&RUN). **D-E)** Heatmaps of LSD1 and histone marks CUT&RUN signal before/after treatment at Corin-activated peaks sorted by LSD1 occupancy for **D)** H3K4me2, H3K27ac, and **E)** H3K4me3 following combination treatment with ATRA. **F)** Distribution of significant Retinoic Acid Receptor (RAR) binding motifs within Corin-activated genomic regions. **G)** Motif analysis of Corin-specific peaks identified retinoic acid receptor (RAR) binding motifs in 51.6% of peaks.

To better understand localized changes in H3K4 methylation and H3K27 acetylation we employed multiplex middle-down proteomics to quantify how drug treatment changes co-modifications of H3. Treatment with either Corin alone or in combination led to acetylation of histones with repressive methylations at H3K9 and/or H3K27, apparent as depletion of peptides modified with both H3K9me2/3 and H3K27me2/3, and increased peptides modified with H3K9me1/2/3 and/or H3K27me1/2/3 and 2-3 acetylations **(Fig. S6-S9)**. Increased acetylation of histones marked by repressive methylations is consistent with partial de-repression of heterochromatic regions, and thus where differences in H3K4 methylation were expected. Methylation of H3K4 was predominantly detected within this set of multiply acetylated, H3K9/K27 methylated peptides, with Corin alone increasing co-modification with H3K4me1, and combination treatment increasing co-modification with H3K4me3 **(Table S3)**.

Interestingly, a motif analysis revealed more than half of the Corin-specific peaks contained significant retinoic acid receptor (RAR) binding motifs (RARA/B/G) **(Fig. 4G)** as defined in Jaspar2024 [25]. Given the presence of RAR elements at Corin-specific peaks **(Fig. 4F)**, we hypothesize that these genes, despite not showing changes in expression with Corin alone, are likely to be induced upon combination treatment with ATRA.

## DISCUSSION

Although retinoic acid (RA) therapy is highly effective in acute promyelocytic leukemia (APL), its therapeutic potential in acute myeloid leukemia (AML) remains significantly limited. The effectiveness of RA is intricately controlled through a network of co-repressors and co-activators whose dynamic interplay modulates transcriptional outcomes. Our findings add substantial clarity to this regulatory complexity by highlighting a direct physical interaction between retinoic acid receptors (RARα and RARβ) and components of the CoREST complex, particularly LSD1 and HDAC1/2. Thus, in AML, CoREST and p300 compete for binding to both RAR and surrounding chromatin. The deacetylation induced by an enzymatically active CoREST complex helps exclude p300 by removing lysine acetylations bound by the p300 bromodomain (PMID: 30924641), and antagonizing acetylation-dependent K4 methyltransferases (PMID: 38960040). CoREST demethylase activity contributes to a repressive RAR program by excluding remodelers (PMID: 38319148), inhibiting H3K9 and H3K27 demethylases (PMID: 20208542; PMID: 38448585), and directly recruiting repressive transcription factors [54].

Under normal conditions, the CoREST complex, characterized by LSD1’s demethylase activity and HDAC1’s histone deacetylation function, maintains a repressive chromatin state at differentiation-promoting genes. This repression significantly restricts chromatin accessibility and the recruitment of transcriptional coactivators essential for gene activation. Interestingly, research by Maiques-Diaz and colleagues revealed that disrupting LSD1’s interaction with GFI1 could reverse this repression, activate key enhancers, and drive differentiation in AML cells. Their findings underscore the therapeutic promise of targeting the CoREST complex to treat leukemia [26]. In our study, the inhibition of LSD1/HDAC using the dual inhibitor Corin when combined with ATRA, altered the dynamics of activating histone marks and the recruitment of the coactivator p300. Corin treatment alone increases H3K4me1/2 and H3K27ac, while co-administration with ATRA promotes the recruitment of P300 to RAR-bound regulatory regions. This shift was accompanied by a pronounced enrichment of active chromatin marks, notably H3K27ac and H3K4me3, at promoters of classic differentiation markers such as *ITGAM* and *CD86*, along with increased chromatin accessibility at RAR motifs. The observed recruitment of coactivators and resultant shift from repressive to active chromatin modifications aligns with previous reports suggesting that the displacement of repressive complexes is critical for enhancer activation, gene transcription, and subsequent differentiation in AML [26, 27]. Our data specifically illustrates the role of ATRA-Corin combination in enhancing coactivator occupancy and activating histone marks, consistent with literature indicating that coactivator binding and histone acetylation are essential steps in enhancer activation and transcription initiation mediated by RARs.

Corin-specific peaks are predominantly located at intron and intergenic pointing to regulatory effect of Corin at putative enhancer regions. These results align with those by Anastas et al. where they find the same bias for activated regions upon Corin treatment in DIPG [21]. Given that Corin-induced accessibility occurs mainly at enhancer loci, it may also influence alternative splicing as suggested by recent findings [28]. ATRA-Corin treatment not only upregulated key myeloid differentiation-associated genes but also induced genome-wide broadening of H3K4me2 peaks. These extensive histone modification domains suggest enhanced transcriptional activity at genes critical for differentiation, consistent with previous reports linking broad H3K4me3 domains to tumor suppressor genes and genes essential for lineage specification [29] . While earlier studies have associated broader H3K4me3 domains with cell identity and transcriptional stability [30] , our findings raise the possibility that H3K4me2 may serve a similar regulatory role. This coordinated epigenetic remodeling may contribute to the stabilization of cell fate decisions during differentiation, supporting the notion that H3K4me2 broadening represents an additional layer of regulatory control in lineage commitment.

Additionally, the increased occupancy of coactivators like p300 following ATRA-Corin treatment suggests a fundamental shift from poised or repressed enhancers toward fully active enhancer states. This shift was well-evidenced in our middle-down proteomic data by the co-accumulation of activating H3 acetylations and K4 methylation on histones previously marked only by repressive K9 and K27 methylations. These newly active enhancers facilitate transcriptional initiation and elongation, promoting the expression of differentiation-associated genes. This mechanism parallels literature reports that coactivator binding, coupled with histone acetylation, contributes directly to chromatin remodeling and gene activation [31]. Collectively, our results underscore a previously underappreciated mechanism whereby LSD1 and the CoREST complex repress RA-driven transcription by outcompeting coactivator binding, which then induces suppression of differentiation-associated genes. By shifting the RAR complex equilibrium in favor of coactivator-binding, ATRA-Corin enhances chromatin accessibility, and robustly promotes transcriptional activation and differentiation. Thus, our findings significantly enhance the understanding of CoREST inhibitor efficacy in combination with RA treatment, revealing new therapeutic avenues for a broader subset of AML patients beyond those traditionally responsive to retinoids. This approach has strong translational potential to alter the AML therapeutic landscape, including with combination or sequencing approaches that support normal myeloid differentiation and tonal eradication of leukemic clones after cytotoxic or Venetoclax-based therapy, or in combination with other epigenetically targeted differentiation agents (e.g., Menin, FLT3, or IDH inhibitors) in non-APL AML.

## MATERIALS AND METHODS

### Cell lines and cell culture

The human AML cell lines U-937 (RRID:CVCL_0007), MOLM-13 (RRID:CVCL_2119), OCI-AML3 (RRID:CVCL_1844), KASUMI-1 (RRID:CVCL_0589), MV4-11 (RRID:CVCL_0064) and SKNO-1 (RRID:CVCL_2196), were cultured in RPMI1640 medium, 10% heat-inactivated FBS (Sigma-Aldrich), 2% penicillin/streptomycin (Gibco, Thermo Fisher Scientific), and SKNO-1 supplemented with 10 ng/mL GM-CSF (Sigma-Aldrich). Cell lines were maintained in Mycoplasma-free conditions and routinely tested for infection (MycoAlert, Lonza).

### Generation of Stable LSD1 shRNA-Expressing Cell Line

To generate stable shRNA knockdown cells, 2 × 10⁶ HEK293T cells (DSMZ Cat# ACC-305, RRID:CVCL_0045) were seeded in a 10 cm dish and transfected 24 hours later using the calcium phosphate method. The transfection mixture included 8 µg of pLKO.1-shRNA plasmid targeting LSD1 (TRCN0000382379), 2 µg of pCMV-VSV-G, and 6 µg of pCMV-dR8.91 packaging plasmid. After 72 hours, viral supernatants were harvested, filtered through a 0.45 µm polyethersulfone membrane, and used to transduce target cells in the presence of 8 µg/mL polybrene (Millipore-Sigma, #TR-1003-G). Following transduction, cells were cultured in complete medium and selected with puromycin (2 µg/mL; BioGems, #5855822) beginning 24 hours post-infection.

### Western blot

Human AML cells were lysed using RIPA buffer supplemented with protease and phosphatase inhibitors. Protein concentrations were determined using the bicinchoninic acid (BCA) assay. Equal amounts of protein were resolved by SDS-PAGE and transferred to nitrocellulose membranes. After blocking with 5% BSA, membranes were incubated overnight at 4 °C with the appropriate primary antibodies: anti-FLAG (Thermo Fisher Scientific Cat# MA1-91878-D680, RRID:AB_2537624), anti-GAPDH (Abcam Cat# ab8245, RRID:AB_2107448), anti Kat3b/p300 (Abcam Cat# ab275378, RRID:AB_2935873), LSD1 (Abcam Cat# ab17721, RRID:AB_443964), HDAC2 (Abcam Cat# ab12169, RRID:AB_2118547), CoREST (Millipore Cat# 07-455, RRID:AB_310629). Following three washes with TBST, membranes were incubated for 1 hour at room temperature with IRDye-conjugated secondary antibodies using Odyssey CLx (RRID:SCR_014579). Protein signals were then visualized using the Odyssey imaging system using Image Studio Lite (RRID:SCR_013715).

### IC_50_ experiments

Cells were seeded at a density of 2 × 10⁴ cells/mL in 96-well plates (Greiner Bio-One, Cat# 655180). Treatments included DMSO (vehicle control) and serial dilutions of the test compounds. Each treatment condition was performed with five technical replicates per concentration, and the entire experiment was repeated in three independent biological replicates.

### Apoptosis analysis

The AML cells were seeded (2 × 10^5^ cells/mL) and treated with ATRA and Corin at the final concentrations of 0.1 μM for 48 h. Then, the AML cells were harvested and stained with Annexin V-PE and 7-AAD according to BD’s instructions (BD, USA), followed by apoptosis analysis by flow cytometry.

### Statistical analysis of drug combination assay and synergy score analysis

Comparisons of cell viability and differentiation between ATRA-Corin combination treatments and individual treatments (ATRA or Corin alone) were analyzed using one-way ANOVA, with a significance threshold of P < 0.001. Multiple comparisons were assessed using Tukey’s post hoc test (P < 0.001). Results from cell line experiments are expressed as mean ± standard deviation (SD) based on three independent biological replicates. Drug synergy assays were conducted as previously described [32]. Briefly, cells were plated into 96-well plates and treated with varying concentrations of each inhibitor, administered either individually or in combination. Cell viability was measured using the Cell-Titer-Glo assay (Promega) and normalized to DMSO-treated controls to determine the inhibitory response. Synergy analysis, including the generation of two-dimensional (2D) synergy matrices, was performed using the SynergyFinder web application [33] (https://synergyfinder.fimm.fi). Heatmaps were produced to visualize interaction effects, with synergy scores interpreted as follows: scores below −10 indicated antagonism, scores between −10 and 10 reflected additive effects, and scores above 10 indicated synergistic interactions.

### Immunoprecipitation (IP)

IP experiments were done using nuclear fraction of cells. Briefly, cells were lysed on ice for 10 min with Buffer A (10 mM HEPES, 1.5 mM MgCl2, 10 mM KCl, 0.5 mM DTT, 0.05% NP40, pH 7.9) followed by centrifugation at 2,300g for 5 min. Supernatant was discarded and the procedure was repeated. The cell pellet was dissolved in IP300 (half of the volume used for Buffer A) and processed as described in the WB section. 1mg of protein was used for each IP and 500*μ*g of nuclear extract as the input material. IP samples were incubated overnight with 10*μ*g of antibody or IgG followed by 30*μ*l of protein A/G agarose bead slurry (Santa Cruz #sc-2003) for 2h. In the IP experiments performed in the presence of Corin, cells were incubated for 30 minutes at 4 °C in Buffer A containing 1*μ*M of the inhibitors. This concentration was maintained during next steps. Similarly, 10*μ*g/ml ethidium bromide (Sigma-Aldrich, E1510) were used to address DNA-independent protein association. IP material was washed 3× with high-salt buffer and eluted with Laemmli buffer, then loaded for SDS-PAGE.

### Affinity Purification of Flag-RARα, Flag-RARβ, and Flag-Mock Complexes

HEK293T cell lines (DSMZ Cat# ACC-305, RRID:CVCL_0045) stably expressing Flag-tagged RARα, RARβ, or Flag-mock controls were cultured in DMEM supplemented with 10% FBS, Puromycin, and Normocin. For each purification, sixty 15-cm² dishes of cells were harvested. Following cytoplasmic fractionation, nuclear extracts were prepared using 10 mL of extraction buffer containing 1.5 mM MgCl₂, 20 mM Tris-HCl (pH 7.9), 0.5 mM DTT, 0.42 M NaCl, 25% glycerol, 0.2 mM EDTA, and 0.2 mM PMSF. Nuclear lysates were dialyzed overnight at 4 °C in 4 L of BC150 buffer (20 mM Tris-HCl pH 7.6, 0.2 mM EDTA, 10% glycerol, 10 mM 2-mercaptoethanol, 0.2 mM PMSF, and 150 mM KCl). The Flag-tagged complexes were purified using 1 mL of anti-FLAG M2 affinity gel (Sigma-Aldrich, A2220) by overnight incubation at 4 °C. The affinity matrix was washed twice with 14 mL of BC500 buffer (20 mM Tris-HCl pH 7.6, 10% glycerol, 0.2 mM EDTA, 10 mM 2-mercaptoethanol, 0.2 mM PMSF, and 0.5 M KCl), followed by three washes with 14 mL of BC150 buffer for 10 minutes each. Bound protein complexes were eluted using 500 µL of 0.5 µg/µL FLAG peptide (Sigma-Aldrich, F3290), prepared according to the manufacturer’s instructions.

### Co-Immunoprecipitation (Co-IP)

Cells were gently washed twice with ice-cold PBS before adding 500 µL of ice-cold IP lysis buffer (25 mM Tris-HCl pH 7.4, 150 mM NaCl, 1% NP-40, 1 mM EDTA, 5% glycerol), supplemented with protease and phosphatase inhibitors (50 µL Halt Inhibitor Cocktail, ThermoFisher). The cells were incubated on ice for 20 minutes with occasional mixing. Lysates were transferred to microcentrifuge tubes and clarified by centrifugation at ∼13,000 × g for 10 minutes at 4 °C. The supernatant was collected, and protein concentrations were quantified using the Pierce BCA Protein Assay Kit (Thermo Scientific, 23250). For each immunoprecipitation, 20 µL of protein A/G magnetic beads (ThermoFisher, 26126, 0.4 µg/µL binding capacity) were washed twice with 500 µL lysis buffer and resuspended in 100 µL lysis buffer containing inhibitors. A/G beads were incubated with 10 µg of antibody, RARβ (Abcam Cat# ab53161, RRID:AB_882283), KDM1A (Abcam Cat# ab17721, RRID:AB_443964), or IgG control (Thermo Fisher Scientific Cat# 02-6102, RRID:AB_2532938), at room temperature for 1 hour with rotation, then continued for an additional hour at 4 °C. After antibody coupling, beads were washed once with lysis buffer. Next, 2 mg of protein lysate was added to the antibody-bound beads and incubated overnight at 4 °C with rotation. The following day, beads were washed twice with 500 µL lysis buffer. For elution, 42 µL of 1x protein sample loading buffer (LI-COR, 928-40004, diluted in lysis buffer) was added, and samples were heated at 90 °C for 10 minutes. 500*μ*g of nuclear extract was used as the input material and twenty microliters of the eluate were loaded onto Bio-Rad 4-15% TGX precast gels for subsequent analysis.

### IP Mass Spectrometry (MS) Analysis

Following immunoprecipitation of Flag-RARα, Flag-RARβ, or Flag-mock controls in HEK293T cells, 50 µL of the final eluates were submitted for mass spectrometry analysis (n=2 replicates per condition). Samples were digested in-solution with trypsin and purified using C18 spin columns. Peptides were then analyzed using a 1.5-hour liquid chromatography gradient on either a Thermo Scientific Q Exactive HF Orbitrap LC-MS/MS System (RRID:SCR_020545) or Thermo Q Exactive HF Hybrid Quadrupole-Orbitrap Mass Spectrometer (RRID:SCR_020558). MS data were searched with full tryptic specificity against the UniProt Human database (07/21/2022, [34]) using MaxQuant 1.6.17.0. [35]. MS data were also scanned for the common protein N-terminal acetylation, Met oxidation and Asn deamidation. Protein and peptide false discovery rates were set at 1%. Label-free quantification (LFQ) normalization was calculated using the MaxLFQ algorithm [36]. Proteomic data were analyzed using the Perseus computational platform (Tyanova, Temu, et al., 2016), following standard procedures for label-free interaction analysis. Reverse hits, proteins identified by site only, and common contaminants were excluded. Intensity Based Absolute Quantification (iBAQ) intensities were log2-transformed, and proteins were retained for analysis if they had at least two non-zero values in either the IP or mock groups (n=2 per group). Missing values were imputed using a normal distribution (width = 0.3, downshift = 1.8). Statistically significant interactors were identified using a two-sample t-test with an s₀ parameter of 2 and a permutation-based FDR threshold of <0.001.

### Silver Staining

Silver staining was performed as previously described [37], with minor modifications. Briefly, protein samples were separated on a 4–20% Tris-glycine gel (Invitrogen) and fixed in a solution containing 10% acetic acid and 50% methanol for 1 hour at room temperature. For further fixation, the gel was incubated in 10% methanol and 7% acetic acid for an additional hour. The gel was then washed with 10% glutaraldehyde for 15 minutes, followed by three 15-minute washes in Milli-Q water. Staining was carried out for 15 minutes using 100 mL of freshly prepared staining solution containing 1 g AgNO₃, 2.8 mL NH₄OH, 185 µL NaOH (10 N stock), and brought to volume with Milli-Q water. After three quick 2-minute washes in Milli-Q water, the gel was developed in 100 mL of developing solution composed of 52 µL 37% formaldehyde and 0.5 mL of 1% citric acid in Milli-Q water. The staining reaction was stopped with 100 mL of stop solution containing 50% methanol and 5% acetic acid.

### Chromatin Immunoprecipitation followed by Sequencing (ChIP-seq) and Library Preparation

AML cells were washed twice with ice-cold PBS and crosslinked in 1% freshly prepared formaldehyde for 10 minutes at room temperature with rotation. Crosslinking was quenched by adding 2.5 M glycine for 5 minutes. Cells were washed with PBS, scraped on ice, pelleted at 600 × g for 10 minutes at 4°C, and washed again twice with cold PBS. Cell pellets were flash-frozen in liquid nitrogen and stored at -80°C. For chromatin preparation, frozen pellets were thawed and resuspended in Lysis Buffer 1 (50 mM HEPES-KOH, pH 7.5; 140 mM NaCl; 1 mM EDTA; 10% glycerol; 0.5% NP-40; 0.25% Triton X-100), incubated on a rotator for 10 minutes at 4°C, and centrifuged at 600 × g for 10 minutes. Pellets were sequentially washed with Lysis Buffer 2 (10 mM Tris-HCl, pH 8.0; 200 mM NaCl; 1 mM EDTA; 0.5 mM EGTA) and Lysis Buffer 3 (10 mM Tris-HCl, pH 8.0; 100 mM NaCl; 1 mM EDTA; 0.5 mM EGTA; 0.1% sodium deoxycholate; 0.5% N-lauroylsarcosine). Chromatin was sonicated in 15 mL tubes (Diagenode Cat. #C01020031) using a Bioruptor Pico for 20 minutes (30 sec ON/OFF cycles) in the presence of 1.2 g of sonication beads at 4°C. Sonicated lysates were centrifuged at 16,000x g for 15 minutes at 4°C to collect chromatin. To assess sonication efficiency, 50 µL of lysate was incubated at 65°C for 3 hours to reverse crosslinks. DNA was then purified using the DNA Clean & Concentrator-5 kit (Zymo Research) and analyzed on a 1% agarose gel. Protein concentration of the remaining lysate was quantified using the Pierce BCA Protein Assay Kit (Thermo Scientific, Cat. #23250). For immunoprecipitation, 2 µg of antibody (H3K4me2, Abcam Cat. #ab32356 or rabbit IgG control, Invitrogen Cat. #02-6102) was pre-bound to 20 µL of Dynabeads Protein A/G (ChIP-grade) in PBS/0.5% BSA overnight at 4°C. Beads were washed, and ∼1–2 mg of chromatin was incubated with antibody-bound beads overnight at 4°C with rotation. Beads were sequentially washed six times with high-salt Wash Buffer (50 mM HEPES-KOH, pH 7.6; 500 mM LiCl; 1 mM EDTA; 1% NP-40; 0.5% sodium deoxycholate) and once with TE buffer containing 50 mM NaCl. Immunocomplexes were eluted by incubating beads in 210 µL Elution Buffer (50 mM Tris-HCl, pH 8.0; 10 mM EDTA; 1% SDS) at 65°C for 15 minutes with shaking (1,000 rpm). Inputs (1%) were processed in parallel. Supernatants were collected and reverse crosslinked overnight at 65°C. The next day, samples were treated with RNase A (1mg/mL; Invitrogen, #12091-021) for 40 minutes at 37°C, followed by Proteinase K (1 mg/mL; Invitrogen Cat. #AM2548) for 3 hours at 55°C. DNA was purified using phenol-chloroform extraction and eluted in 50 µL nuclease-free water. DNA concentrations were measured using a Qubit 3 fluorometer with dsDNA High Sensitivity reagents (Thermo Fisher Scientific, Cat. #Q32851). ChIP DNA was used for library preparation with the NEBNext Ultra DNA Library Prep Kit for Illumina (New England Biolabs Cat. #E7370) according to the manufacturer’s instructions. Fragment size distribution was analyzed using a 2200 TapeStation Instrument (RRID:SCR_014994) with D1000 ScreenTape (Agilent Technologies, Cat. #5067–5582). Libraries were quantified with Qubit and sequenced (generating a minimum of 30 million single-end reads per sample, read length: 100 bp) on the Illumina NovaSeq 6000 Sequencing System (RRID:SCR_016387).

### ChIP-seq Analysis

ChIP-seq data were processed with the standardized ChIP-seq ENCODE Pipeline (v2) (https://github.com/ENCODE-DCC/chip-seq-pipeline2) [38]. Raw FASTQ files were processed with Trimmomatic (v0.39) [39]. Trimmed reads were aligned to the human reference genome (hg38) using Bowtie2 (v2.2.6.) [40]. Duplicate reads were removed using Picard (v1.126) [41], and alignments were filtered to retain uniquely mapping reads with a mapping quality of ≥30. Upon discarding annotated blackregions [42], deepTools2 [43] was used to generate CPM normalized bigwig (bamCoverage), and heatmaps (computeMatrix and plotHeatmap functions). Peaks were identified using MACS2 [44] with default parameters for narrow peaks and additional parameters (--broad, --broad-cutoff 0.1) for histone modifications showing broader enrichment patterns. Irreproducible Discovery Rate (IDR) analysis (v. 2.0.4) [45] was employed to evaluate peak reproducibility across replicates generating a high confidence set of peaks. The peaks significantly enriched compared to their corresponding Input control for a given treatment (optimal peaks with fold change > 3 and a q-value < 0.01) were used for downstream analysis. HOMER (annotatePeaks function) [46] was used for peak annotation. BEDTools [47] function intersect was used to generate common sets of peaks where needed. Peak coverage per gene was performed with an in-house python script using all significant peaks annotated to genes.

### CUT&RUN Assay and Sequencing

CUT&RUN was performed using the CUTANA™ ChIC/CUT&RUN Kit (EpiCypher, Cat. No. 14-1048) following the manufacturer’s instructions. Briefly, 500,000 MOLM-13 cells were harvested and assessed for full permeability using digitonin and Trypan blue staining to determine optimal conditions. Permeabilized cells were then bound to activated Concanavalin A beads and incubated overnight at 4 °C with the following primary antibodies: anti-H3K27ac (EpiCypher, SKU: 13-0059), anti-H3K4me2 (EpiCypher, SKU: 13-0027), anti-H3K4me3 (EpiCypher, SKU: 13-0060), anti-IgG control (EpiCypher, SKU: 13-0042K), and anti-LSD1 (Abcam, ab17721). After extensive washing, cells were incubated with pAG-MNase (EpiCypher, SKU: 15-1016) for 10 minutes at room temperature, followed by chromatin digestion for 2 hours at 4 °C. The reaction was stopped by adding stop buffer and E. coli spike-in DNA (EpiCypher, SKU: 18-1401). DNA fragments were then purified using DNA cleanup columns. Libraries were prepared with the CUTANA™ DNA Library Prep Kit (EpiCypher, SKU: 14-1001) and sequenced on an Illumina NovaSeq 6000 Sequencing System (RRID:SCR_016387) generating a minimum of 5 million paired-end reads per sample (read length: 150 bp).

### CUT&RUN Analysis

FASTQ files were processed and analyzed using the NF-core cutandrun pipeline (doi: 10.5281/zenodo.5653535) [48, 49], within Pluto bioinformatics platform (https://pluto.bio). Briefly, paired-end FASTQ files were quality-filtered and trimmed using TrimGalore . Reads were aligned to the human genome GRCh38 reference genome (NCBI build p.14, release 110) using Bowtie2 [39], and duplicate reads were removed from control samples using Picard [41]. Peak calling was performed using MACS2 [44] Peaks were defined as genomic regions where at least one target sample significantly surpassed its corresponding IgG control. Consensus peaks across all replicates of each treatment were merged using BEDTools and annotated to the nearest transcriptional start site (TSS) using HOMER [46]. Peak counts for downstream analyses were generated with featureCounts [50].

### RNA Sequencing (RNA-seq)

For RNA sequencing, 500 ng of total RNA was extracted using TRIzol reagent (Thermo Fisher Scientific, Cat. #15596026) and treated with DNase I (RNase-free, Cat. #AM1907) to remove any residual genomic DNA. Libraries were prepared using the TruSeq Stranded Total RNA Library Prep Kit (Illumina, Cat. #20020596) following the manufacturer’s protocol. Sequencing was performed on the Illumina NovaSeq 6000 Sequencing System (RRID:SCR_016387), generating a minimum of 50 million paired-end reads per sample (read length: 100 bp).

### RNA-seq Analysis

Raw FASTQ files were quality-filtered and trimmed using Trimmomatic (RRID:SCR_011848) [39] to remove adapter sequences and low-quality bases. Trimmed reads were aligned to the human reference genome (hg38) using STAR aligner (RRID:SCR_015899) [51] with default settings. We obtained >40 millions of aligned reads per sample. Gene-level expression quantification was performed using RSEM (RRID:SCR_013027) [52] against the human Ensembl reference transcriptome (release 97), using an expectation-maximization approach. Gene expression levels were normalized to Transcripts Per Million (TPM). Differential expression analysis by DEseq2 was performed at a 2-fold change cutoff and an FDR<0.05 significance cutoff, unless noted otherwise. GSEA enrichment analysis [53] was performed on pre-ranked data obtained from Deseq2 (ranking based on the “stats” column) against Hallmarks of cancer. The ATRA-Corin specific gene signature includes genes differentially expressed by ATRA-corin treatment, and not differentially expressed by neither ATRA nor corin treatments alone. Heatmaps showing Z-score row-normalized gene expression and unsupervised hierarchical clustering were generated using clustermap (method: ward; metric: euclidean distance, https://doi.org/10.21105/joss.03021) to visualize patterns of differential gene expression. Enrichment analysis was performed on the differential expressed genes by Enrichr [54] against GO biological process.

### ATAC-seq and Data Analysis

ATAC-seq was carried out following the general protocol established [55]. DNA concentrations were measured using a Qubit 3 fluorometer with dsDNA High Sensitivity reagents (Thermo Fisher Scientific). Libraries were prepared, pooled, and sequenced on the Illumina NovaSeq 6000 Sequencing System (RRID:SCR_016387) to generate 75 bp paired-end reads.

ATAC-seq data were processed using the ENCODE ATAC-seq pipeline (v1) with default parameters. Paired-end raw FASTQ reads trimmed using cutadapt v1.9.1 [56]. Trimmed reads were aligned to the human reference genome (hg38) using Bowtie2 v2.2.6 [40]. Resulting BAM files were filtered to exclude mitochondrial reads and PCR duplicates using SAMtools v1.7 [57] and Picard MarkDuplicates v1.126 [41]. Paired-end reads were processed to retain only properly paired, uniquely mapped reads with mapping quality ≥30. Reads were shifted +4 bp (forward strand) and −5 bp (reverse strand) to account for Tn5 insertion bias. Peaks were called using MACS2 v2.1.0 [44] (--nomodel --shift 75 --extsize 150) with a p-value threshold of 0.01. Replicate concordance and IDR analysis were performed analogously to ChIP-seq data. Signal tracks were generated for individual replicates and for pooled replicates. Accessibility heatmaps were generated using the pooled signal from 2 replicates per condition using deepTools2 *computeMatrix* and *plotHeatmap* functions.

### Motif analysis

Motif analysis was performed with fimo package within the MEME suite [58] to identify peaks containing RARE motifs in addition to unbiased motif searches using the annotated motifs in JasparDB [59].

### Middle-down MS

MOLM13 cells were treated in triplicate for 6 hours with Corin (100 nM), ATRA (200 nM), Corin and ATRA (100 nM and 200 nM respectively), or DMSO vehicle. After treatment cells were harvested, pelleted, washed and snap frozen for storage at -80 C. Frozen cell pellets (10^7 / pellet) were placed on ice and gently resuspended in 1 mL of ice-cold lysis buffer (10 mM Tris, pH 7.9, 1.5 mM MgCl2, 10 mM KCl, 0.5 mM DTT, 0.2 mM PMSF, 10 mM sodium butyrate), and placed on an end-over-end rotator at 4 C. After 10 minutes the resuspended cells were gently pipetted to break up any remaining clumps, and returned to the end-over-end rotator for a further 10 minutes at 4 C. Nuclei were pelleted by centrifugation (10 minutes, 900 RCF, 4 C) and the supernatant was removed, after which the pelleted nuclei were gently resuspended in 1 mL resuspension buffer (20 mM HEPES, pH 7.8, 25 mM NaCl, 5 mM MgCl2, 0.25 M sucrose, 0.5 mM DTT, 10 mM sodium butyrate) supplemented with protease inhibitor (Pierce Protease Inhibitor Tablets, EDTA-free, Thermo Scientific), and placed on an end-over-end rotator for 5 minutes at 4 C. Nuclei were again pelleted by centrifugation, the supernatant removed, and the pellet gently washed with resuspension buffer as before. After removal of the wash supernatant, the nuclear pellet was vigorously mixed with 0.2 N sulfuric acid (100 uL / pellet) and placed on an end-over-end rotator for 2 hours at 4 C. Nucleic acid and nuclear debris was removed by centrifugation (15 minutes, 10,000 RCF, 4 C) and the supernatant was dialyzed (Slide-a-lyzer mini dialysis cups, 3.5 kDa MWCO, Thermo Scientific) sequentially against 1 L ice-cold 2.5% acetic acid (18 hours, 4 C), 1 L ice-cold 0.05% trifluoroacetic acid (2 hours, 4 C) and 1 L ice-cold 0.05% trifluoroacetic acid (2 hours, 4 C). Protein concentration in the nuclear extract supernatant was estimated by absorbance at 280 nm (NanoDrop2000, Thermo Scientific), after which the supernatant was aliquoted, snap frozen and freeze dried before storing at -80 C. Freeze dried nuclear extracts were resuspended to 15 μM histone H3 in a concentrated protease inhibitor buffer (50 mM PIPES, pH 7.5, 6 mM CaCl2, 3 mM PMSF) and incubated for 10 minutes on ice. Excess PMSF was quenched with DTT (11 mM) and incubated for 10 minutes at 20 C. Tandem mass tag labelled oligoglycine peptide (1 per biological replicate) and cW11 sortase were added and placed in a 37 C incubator for 1 hour (final concentrations: 40 mM PIPES, 5 mM CaCl2, 2.5 mM PMSF, 10 mM DTT, 1 mM oligoglycine peptide, 0.2 mM sortase, 10 μM histone H3). After 1 hour at 37 C additional cW11 sortase was added (final concentration: 0.4 mM cW11) and the reaction was returned to the 37 C incubator for a further 6 hours. Reactions were chilled on ice before adding trichloroacetic acid to a 5% (v/v), then mixed vigorously, and the precipitated protein was pelleted by centrifugation (20 minutes, 13,200 rcf, 4 C). The reaction supernatant, containing labelled histone tail peptides, was bound to stage tips packed in house, and washed with 0.1% trifluoroacetic acid. Peptides were eluted with 80% acetonitrile, 20% water, 0.1% formic acid and freeze dried.

Peptides were resuspended in water and quantified by SDS-PAGE against a peptide standard. Peptides were reconstituted in 0.1% formic acid to 10 ng/uL, and mixed at 1:1:1:1:1:1 mass ratios into 6-plex sets. Samples (20 ng / run) were analyzed with a Vanquish Neo LC (Thermo Fisher Scientific) coupled to an Orbitrap Ascend Tribrid mass spectrometer equipped with ETD, PTCR, and UVPD (Thermo Fisher Scientific). The LC was operated in trap-and-elute mode using a PepMap™ Neo C18, 100 Å, 5 μm, 0.3 x 5 mm Trap Cartridge (Thermo Fisher Scientific) and an Easy-Spray Pep-Map Neo C18, 100 Å, 2 μm, 75 μm x 150 mm column (Thermo Fisher Scientific) heated to 40°C at a flow rate of 0.3 μL/min. The LC gradient was 2 to 15% Solvent B in 40 min, followed by 15 to 25% Solvent B in 6 min (Solvent A: 0.1% formic acid; Solvent B: 0.1% formic acid in acetonitrile). The mass spectrometer was operated in data-dependent acquisition mode with a 2 second cycle time. MS1 spectra were collected in the Orbitrap at a resolution of 60k and over a scan range of 375-600 m/z. Charge state and precursor selection range filters were applied to select peptides with a charge state of 8 from 480-540 m/z for MS2. Dynamic exclusion was enabled, and precursors were excluded for 9 seconds after being selected 1 time. MS2 spectra were collected with an isolation window of 1.6 m/z, a resolution of 30k, and 2 microscans. EThcD was performed with an ETD reaction time of 20 ms, ETD reagent target of 1E5, max ETD reagent injection time of 20 ms, supplemental activation collision energy of 15%, and normalized AGC target of 200%.

### Middle-down MS analysis

Raw files were processed as previously described. Briefly, raw data files were analyzed using Byonic MS/MS search engine (Protein Metrics) and the following settings:No protease cleavage; 500 ppm precursor tolerance; 15 ppm fragment tolerance; maximum 5 precursors; fragmentation type EThcD; maximum 6 common PTMs; Common PTMs Methyl / +14.015650 @ K, R, max 3 | Dimethyl / +28.031300 @ K, R, max 3 | Trimethyl / +42.046950 @ K, max 3 | Phospho / +79.966331 @ S, T, max 2 | Acetyl / +42.010565 @ K, max 5; maximum 1 rare PTM; Propionyl / +56.026215 @ K | rare1; fixed modification of H with TMT 6-plex; TMT6plex / +229.162932 @ H; fixed modification of protein C-terminus with 76.1001 Da mass (accounting for the mass difference between H and [K + 2,3-diaminopropionamide]).

Data was searched against a fasta database of histone protein sequences, common contaminants, and the following sequences for H3.1 and H3.3 peptides:

H3.1 ARTKQTARKSTGGKAPRKQLATKAARKSAPATGGGH

H3.3 ARTKQTARKSTGGKAPRKQLATKAARKSAPSTGGGH

Peptide spectrum matches (PSMs) from Byonic were exported and subsequently processed in R (4.4.2) using RStudio, custom scripts (available upon request) and packages parallel (4.4.2), stats (4.4.2), purrr (1.1.0), mzR (2.40.0), Rcpp (1.1.0), tidyr (1.3.1), stringr (1.5.2), dplyr (1.1.4), xml2 (1.4.0), readr (2.1.5), readxl (1.4.5), writexl (1.5.4), ggplot2 (3.5.2) and ggrepel (0.9.6).

PSMs were filtered for high confidence matches with a posterior error probability ≤ 0.05, and a Byonic delta mod score ≥ 10. Spectra were subdivided by the number of PSMs reported in Byonic, then EThcD ions (a, b, c, y, z, and neutral loss; 15 ppm error tolerance) consistent with the PSM(s) were identified from the spectrum. Fragment ions (excluding TMT ions) with at least one heavy isotope peak (e.g. 15N) were treated as high confidence and used to determine proteoform abundance in the sample. Proteoform abundance was calculated by dividing annotated fragment ion intensity attributable to a single proteoform by the sum of annotated fragment ion intensity for all proteoforms. For chimeric spectra assigned multiple PSMs, intensity from shared fragment ions was divided between the relevant PSMs. Reporter ion signal intensities were channel normalized to the maximum average channel (e.g. TMT-126) signal intensity, averaging over all spectra with a PSM. Channel normalized reporter ion signal intensities were log2 transformed for statistical analysis.

Spectra/scans with ≥2 reporter ions per treatment condition were used in quantification, and all scans attributable to a proteoform (e.g. H3K14ac/K23ac/K27me2) were grouped. For each treatment condition all log2 transformed reporter ion signals from a proteoform grouping were averaged, and the difference between treatment condition averages reported as the log2-fold change in proteoform abundance. Statistical significance was assessed by two-sided T test (p<0.05) using R stats (4.4.2).

## AUTHOR CONTRIBUTIONS

**M.M. Tayari:** Conceived and designed the overall project; led the conceptualization and performed experiments; conducted data analysis, visualization, and methodology; wrote the original draft and contributed to review and editing of the manuscript. **H. Gomes Dos Santos**: performed bioinformatics analyses, data visualization and manuscript review and editing. **S. Dokaneheifard**: Performed FLAG affinity purification, co-immunoprecipitations, silver staining, and western blot analyses. **S.D. Whedon** Performed middle-down proteomics and analysis. **M. Valencia**: Generated FLAG-tagged RARα, RARβ, and mock HEK293T cell lines used in the study. **A. Harikumar:** Performed the ChIP-seq experiments. **F. Beckedorf:** Contributed to bioinformatics analysis, and data visualization. **P. Cole**: Provided project administration support and contributed to manuscript review and editing. **J.M. Watts**: Supported the project through resources, funding acquisition, validation, investigation, and project administration, and participated in reviewing and editing the manuscript. **R. Shiekhattar**: Oversaw project conceptualization, supervision, and funding acquisition, and contributed to validation, investigation, project administration, and final review and editing of the manuscript.

## ACKNOWLEDGEMENTS

This work was supported by funding from the University of Miami Miller School of Medicine, the Sylvester Comprehensive Cancer Center, and the Leukemia & Lymphoma Society’s Specialized Center of Research Program (LLS-SCOR), awarded to Stephen D. Nimer (Program PI), with support for Project 1 to Ramin Shiekhattar, Philip A. Cole, and John M. Watts. Additional support was provided by NIH grant R01GM078455 awarded to R.S., and by the Oncogenomics Shared Resource (OGSR; SCR_022502), which is supported by the National Cancer Institute of the National Institutes of Health under Award Number P30CA240139. The content is solely the responsibility of the authors and does not necessarily represent the official views of the National Institutes of Health. We thank members of the Shiekhattar and Cole laboratories for their valuable discussions and feedback. We are grateful to Arthur Zelent for his early contributions to the project, and to Liliana Garcia-Martinez and Lluis Morey for providing the shLSD1 constructs. Mass spectrometry analyses were performed by the Proteomics and Metabolomics Facility at the Wistar Institute. The graphical abstract was created using BioRender (https://biorender.com).

## DATA AVAILABILITY

All data supporting the findings of this study are available within the main text and/or the Supplementary Materials. Raw and processed RNA-seq, ChIP-seq, ATAC-seq and CUT&RUN datasets have been deposited in the Gene Expression Omnibus (GEO) under accession numbers GSE300986, GSE300987, GSE300988, GSE300989. Raw and processed middle-down proteomics datasets, as well as relevant data processing scripts have been deposited on Harvard Dataverse.

## ETHICS STATEMENT

No human participants or animal studies were involved. Only commercially available and previously established cell lines were used.

