## Supplementary Figures for "Balancing Activation and Repression: CoREST-p300 Antagonism Controls Retinoic Acid-Driven Differentiation in AML"

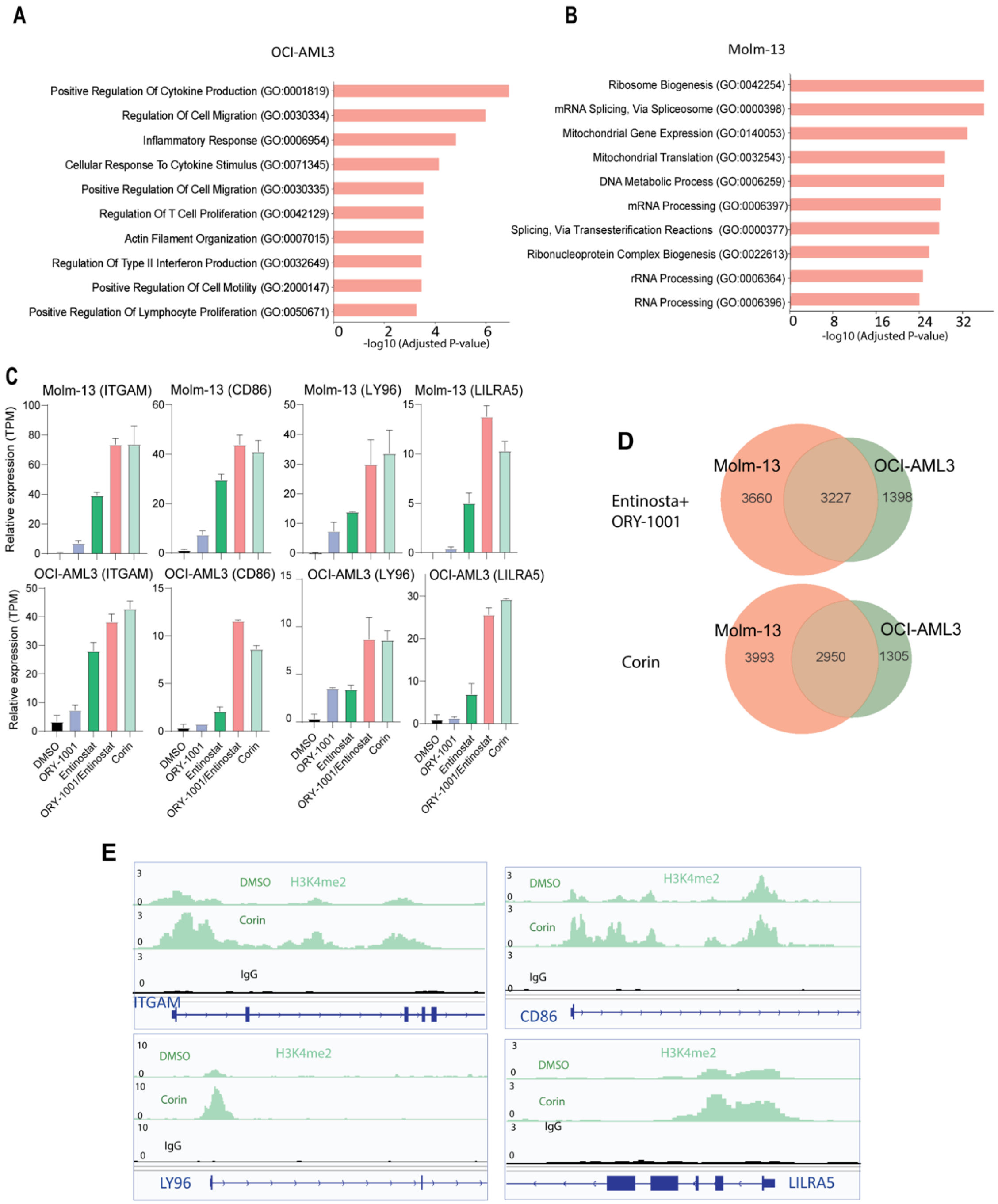


**Figure S1. A)** Gene ontology analysis of genes upregulated in OCI-AML3 and **B)** MOLM-13 cells following Corin treatment. **C)** Relative expression levels (TPM, transcripts per million) of selected differentiation-related genes in MOLM-13 cells (on top) and OCI-AML3 cells (on the bottom). significance here defined by a two-tailed unpaired Student's *t* test. **D)** Venn diagram illustrating gene overlap between OCI-AML3 and MOLM-13 cell lines treated with either ORY-1001 plus Entinostat or Corin alone. **E)** Promoters of differentiation-related genes showing increased H3K4me2 enrichment following Corin treatment. Data analysis and plots were generated using Pluto (https://pluto.bio)

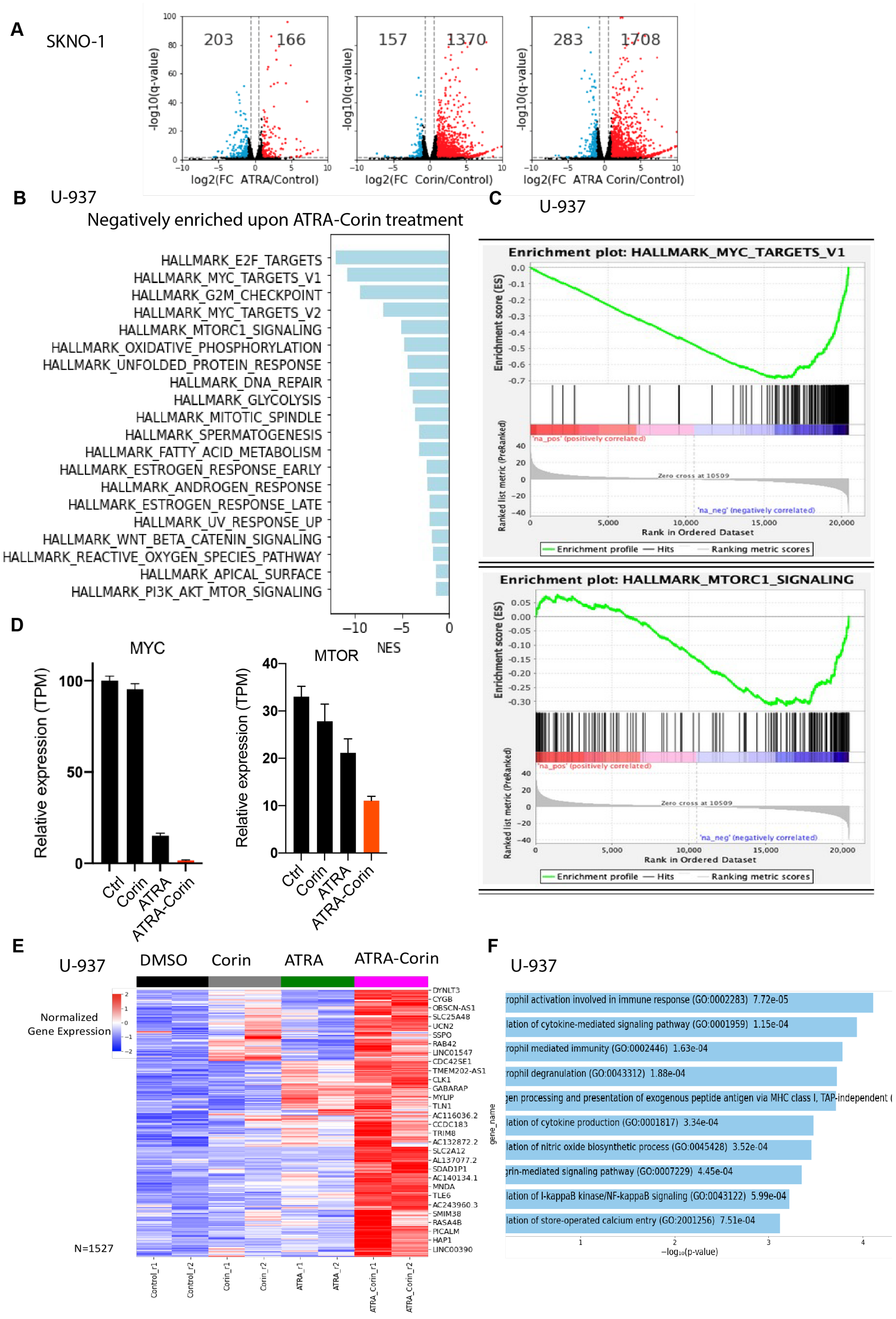


**Figure S2.** Concordant activation of genes involved in hematopoietic lineage commitment, differentiation, and immune response. **A)** Volcano plots showing differential gene expression in SKNO-1 cells treated with ATRA, Corin, or the combination. Genes are color-coded as non-differentially expressed (black), upregulated (red), or downregulated (blue). Cutoffs: FDR ≤ 0.05 and |fold change| ≥ 2. **B)** Gene Ontology (GO) analysis of genes negatively enriched by ATRA-Corin treatment in U-937 cells, but not by either treatment alone. **C)** Enrichment plots showing downregulation of MYC targets and mTORC1 signaling pathways upon ATRA-Corin treatment. **D)** Relative expression levels (transcripts per million, TPM) of *MYC* and *MTOR* genes in U-937 cells.

A

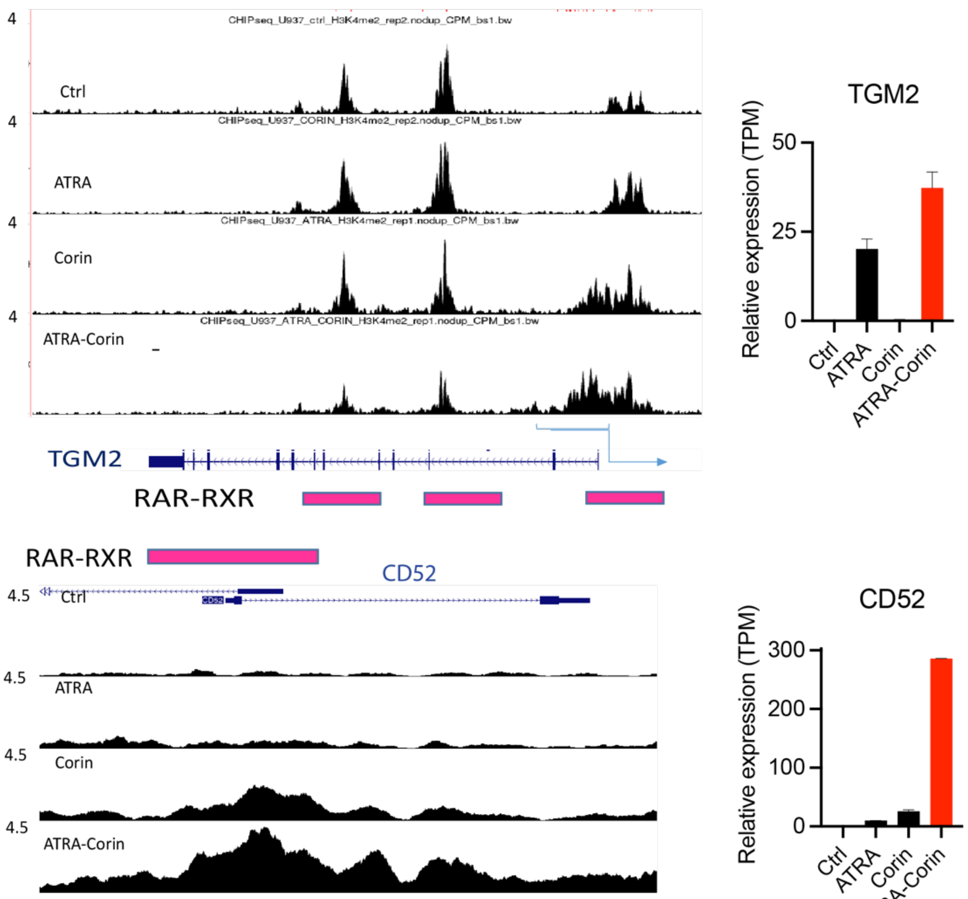


A

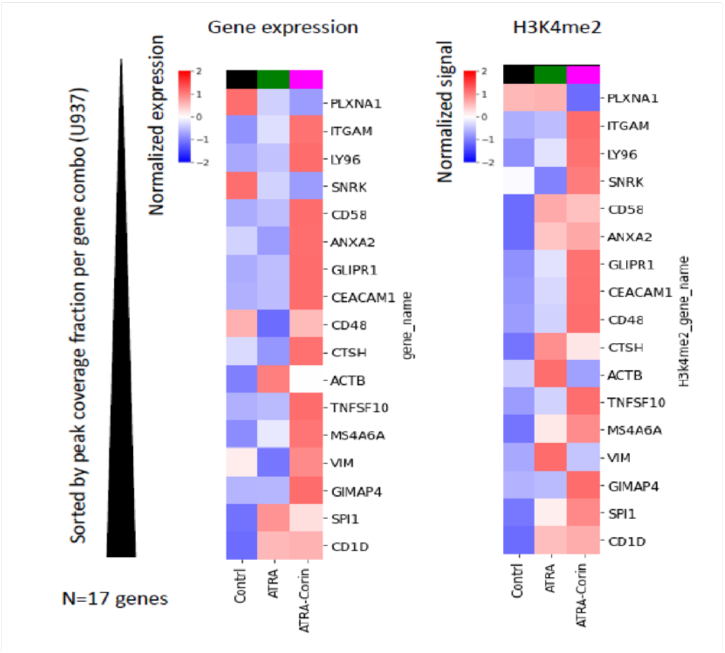


**Figure S3. ATRA-Corin treatment induces broadening of H3K4me2 domains.** Genome-wide analysis revealed a marked expansion of H3K4me2 peaks following treatment.

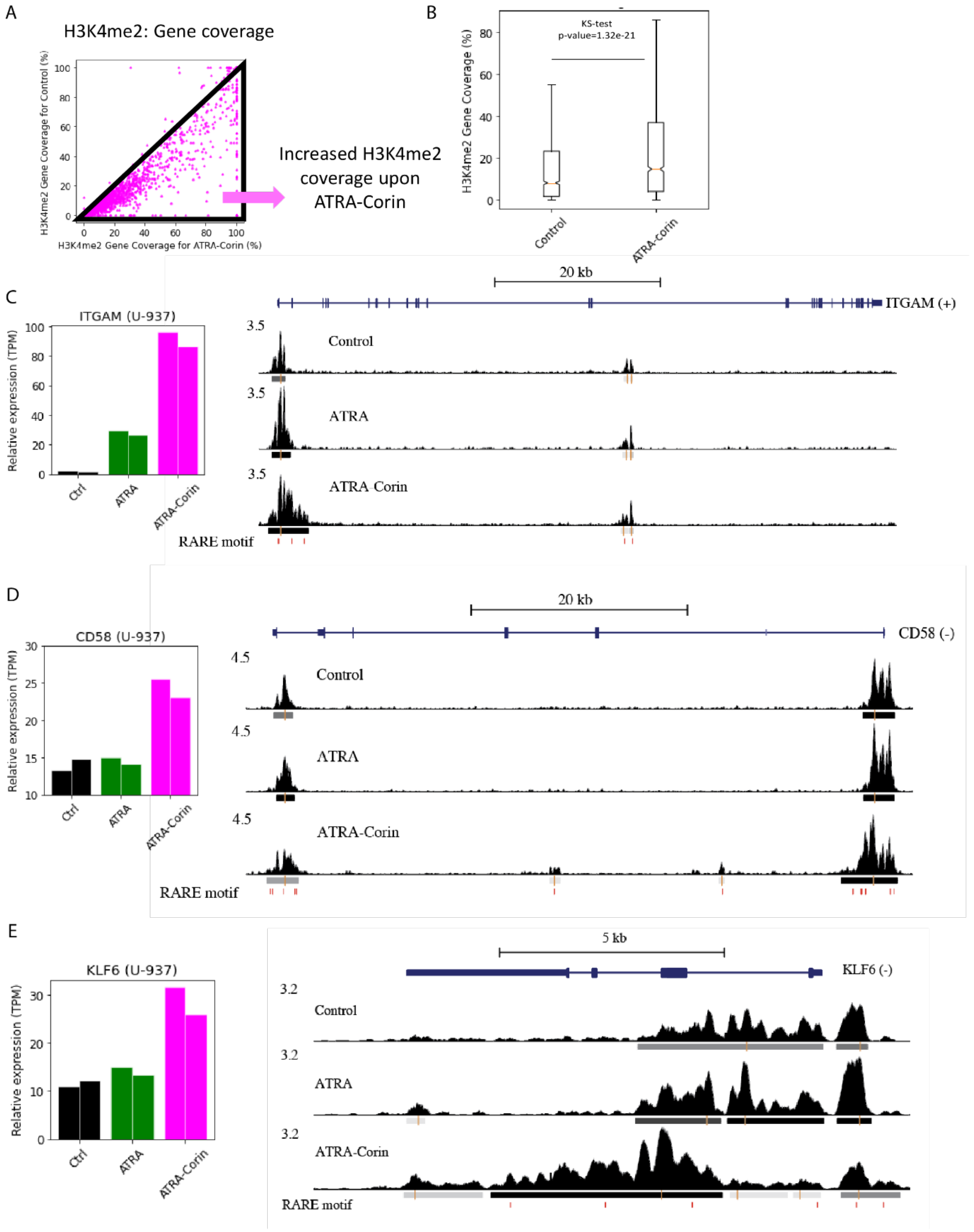


**Supplementary Figure 4.** Changes in H3K4me2 enrichment following ATRA-Corin treatment in U-937 cells. **A)** Genome-wide increase in H3K4me2 coverage after ATRA-Corin treatment. **B)** Percentage of genes showing H3K4me2 enrichment in DMSO-treated vs. ATRA-Corin-treated cells. **C-E)** Relative expression (TPM) of *ITGAM* (C), *CD58* (D), and *KLF6* (E). Retinoic acid response elements (RAREs) are marked by red blocks at the bottom of each panel.

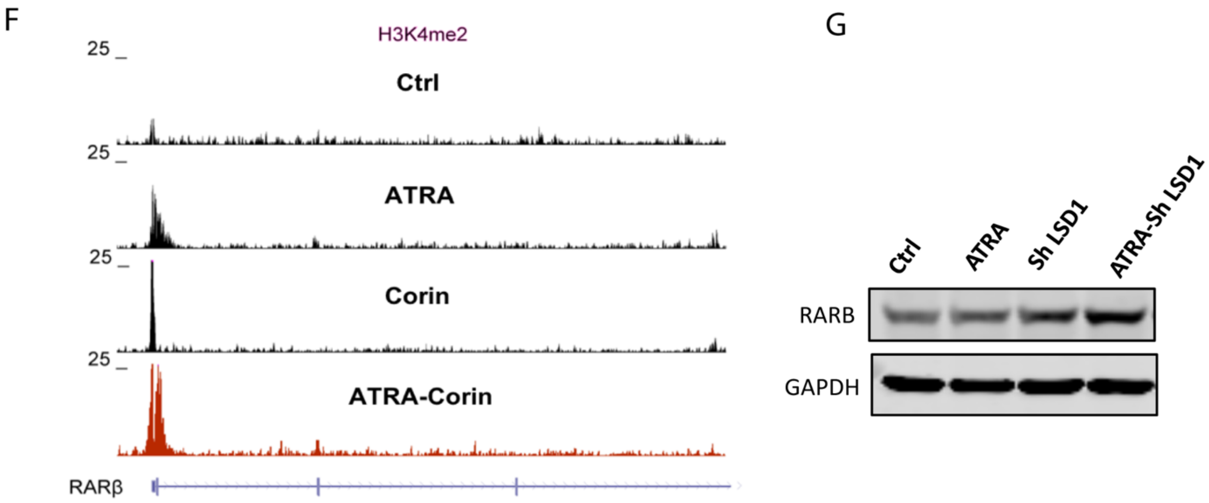


**Figure S4. Regions containing retinoic acid response elements (RAREs; red blocks) show pronounced H3K4me2 broadening, most notably at the *RARB* promoter.** Increased H3K4me2 enrichment at the *RARB* promoter correlates with elevated protein levels. **A)** H3K4me2 levels at the *RARB* promoter region following treatment. **B)** Western blot analysis of RARB protein expression in U-937 cells treated with DMSO, ATRA, Corin, or the ATRA-Corin combination.

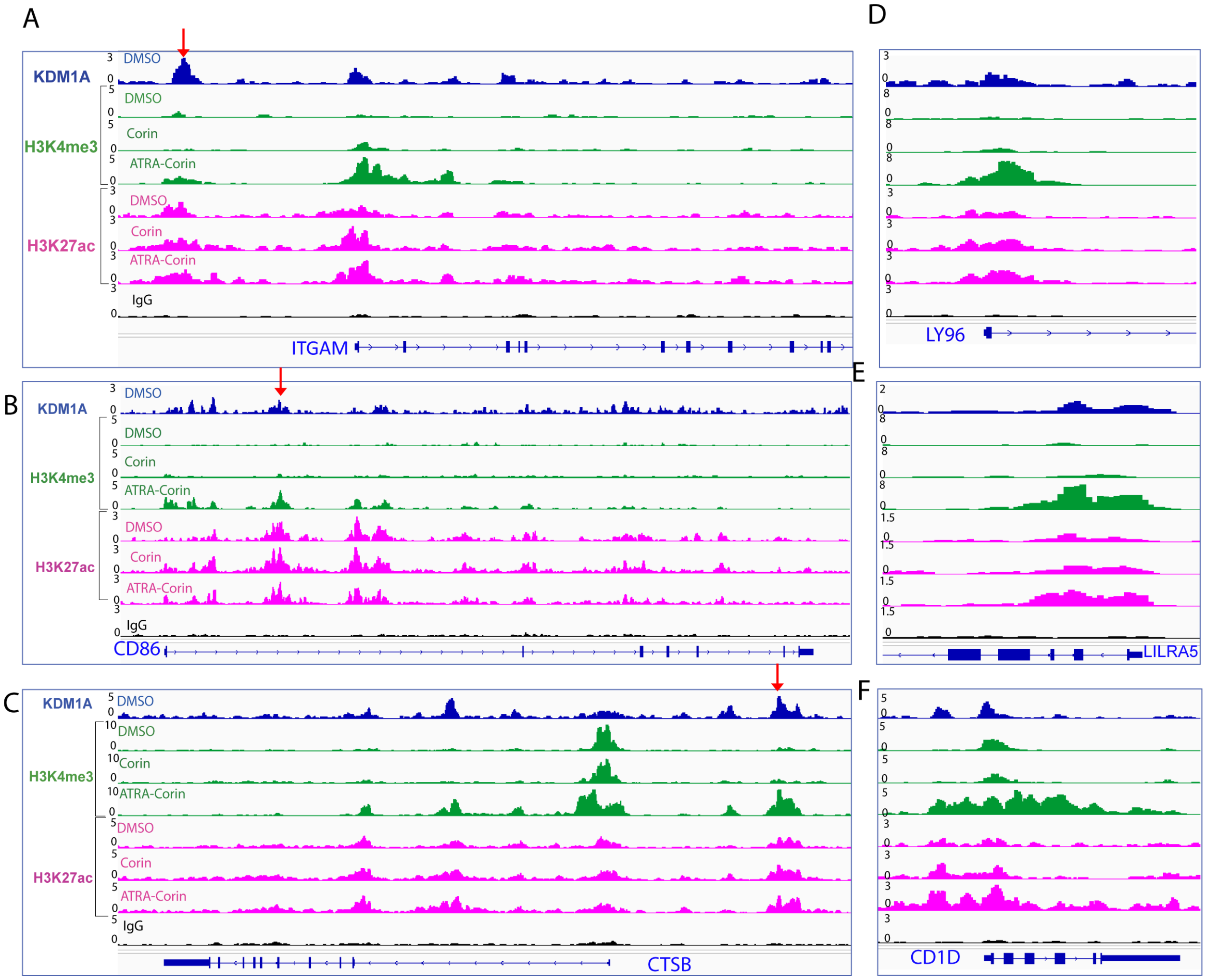


**Figure S5. ATRA–Corin co-treatment enhances activating histone marks at myeloid differentiation loci**. Combined treatment increased H3K4me3 and H3K27ac at target regions *ITGAM*, *CD86*, *CTSB*, *LY96, LILRA5* and *CD1D*. Red arrows highlight enhancers near *ITGAM*, *CD86*, and *CTSB*, showing LSD1 occupancy with increased H3K4me3 and H3K27ac.

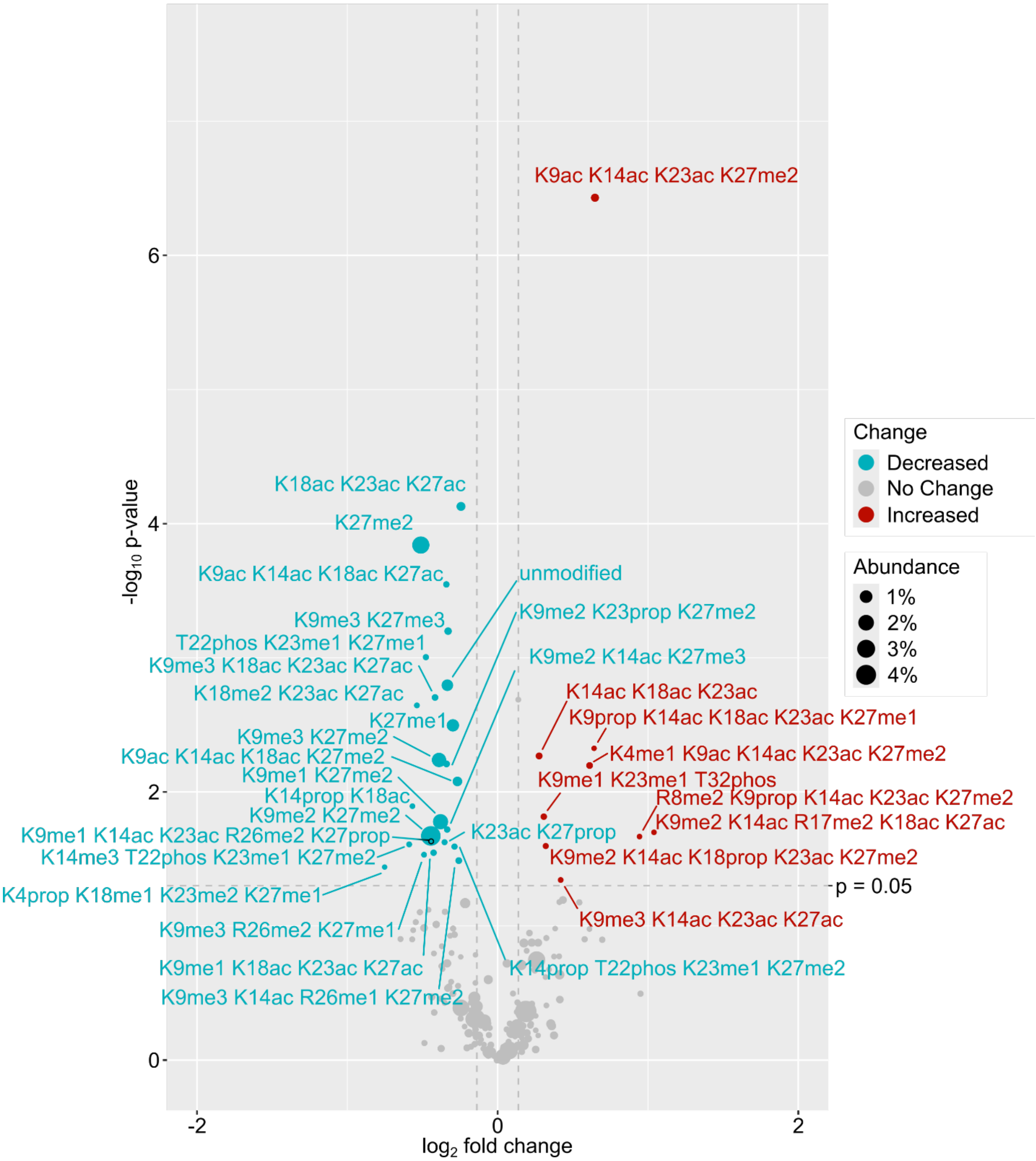


**Figure S6. Combinations of Histone H3 post-translational modifications changed upon treatment with Corin alone compared to DMSO.** Modification patterns significantly increased (red) or decreased (blue) by more than 10% (vertical dashed line) in cells treated with Corin as compared to cells treated with DMSO vehicle. Peptide abundance (relative to all H3 proteoforms detected) is illustrated by point size. Statistical significance of the change ($p < 0.05$) is demarcated by the horizontal dashed line. PTMs: ac = acetyl; me1 = monomethyl; me2 = dimethyl; me3 = trimethyl; prop = propionyl; phos = phosphoryl.

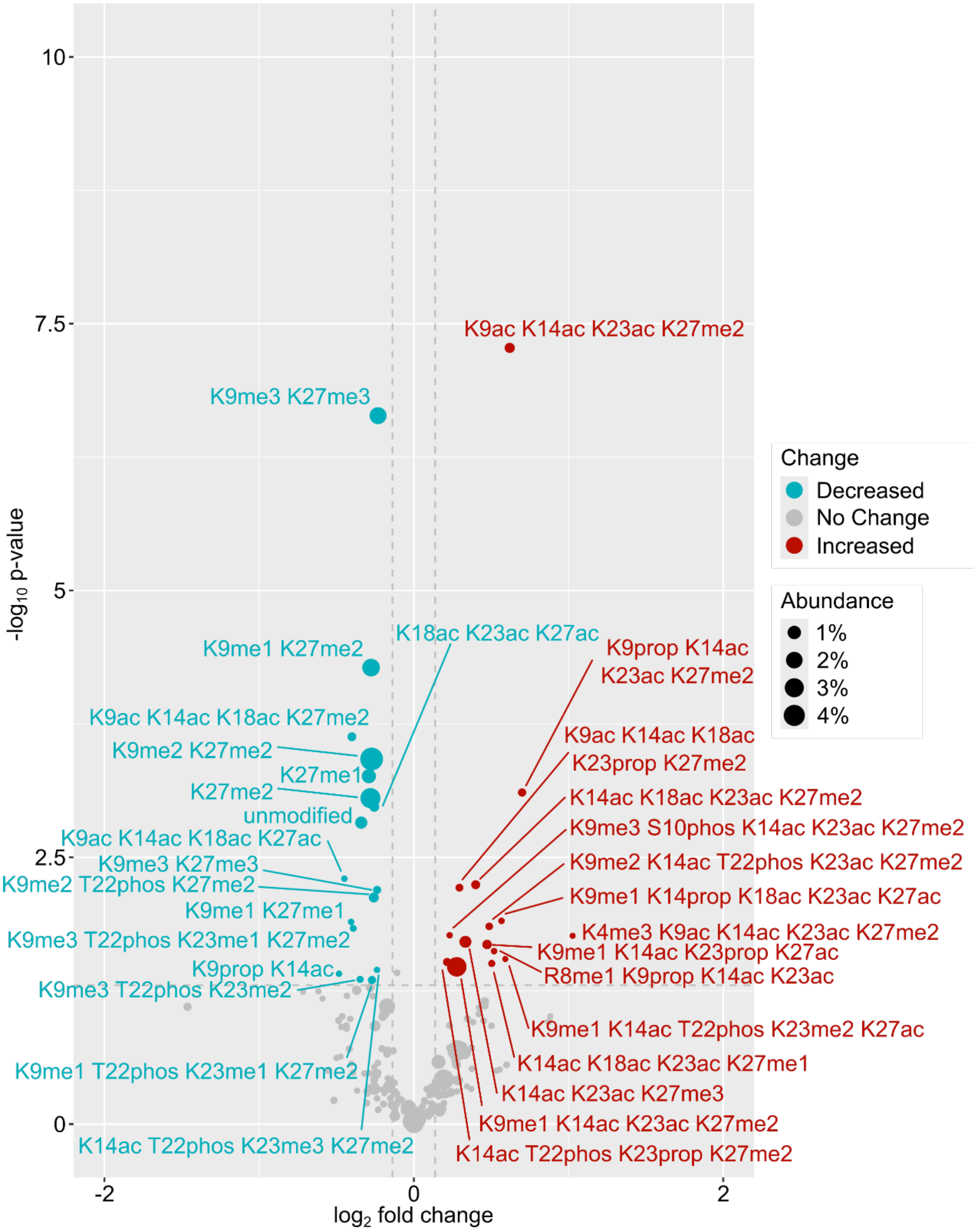


**Figure S7. Combinations of Histone H3 post-translational modifications changed upon combination treatment with Corin and ATRA compared to ATRA alone.** Modification patterns significantly increased (red) or decreased (blue) by more than 10% (vertical dashed line) in cells treated with Corin and ATRA as compared to cells treated with ATRA alone. Peptide abundance (relative to all H3 proteoforms detected) is illustrated by point size. Statistical significance of the change ($p < 0.05$) is demarcated by the horizontal dashed line. PTMs: ac = acetyl; me1 = monomethyl; me2 = dimethyl; me3 = trimethyl; prop = propionyl; phos = phosphoryl.

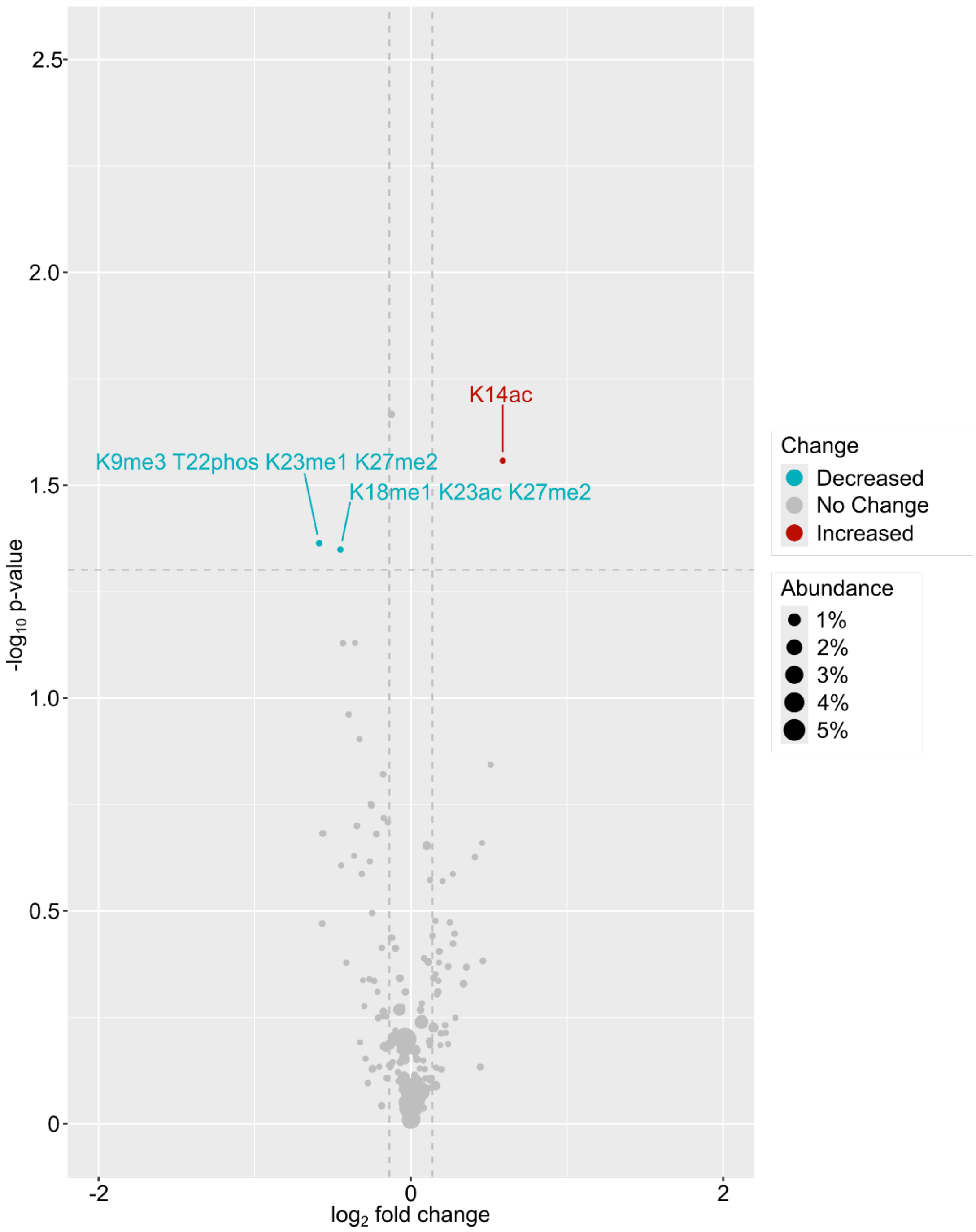


**Figure S8. Combinations of Histone H3 post-translational modifications changed upon treatment with ATRA alone compared to DMSO.** Modification patterns significantly increased (red) or decreased (blue) by more than 10% (vertical dashed line) in cells treated with ATRA as compared to cells treated with DMSO vehicle. Peptide abundance (relative to all H3 proteoforms detected) is illustrated by point size. Statistical significance of the change ($p < 0.05$) is demarcated by the horizontal dashed line. PTMs: ac = acetyl; me1 = monomethyl; me2 = dimethyl; me3 = trimethyl; prop = propionyl; phos = phosphoryl.

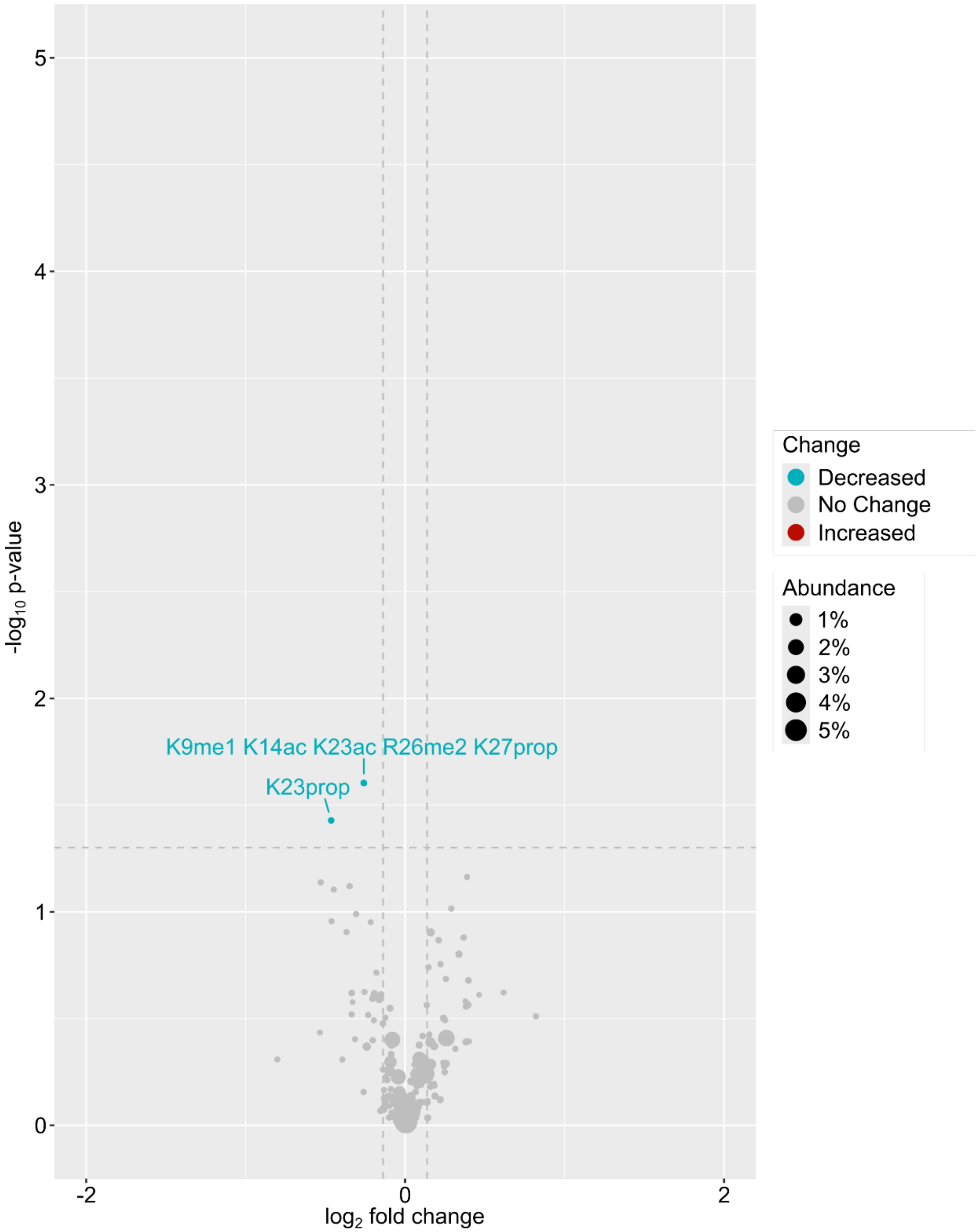


**Figure S9. Combinations of Histone H3 post-translational modifications changed upon treatment with Corin and ATRA compared with Corin alone.** Modification patterns significantly increased (red) or decreased (blue) by more than 10% (vertical dashed line) in cells treated with Corin and ATRA as compared to cells treated with Corin alone. Peptide abundance (relative to all H3 proteoforms detected) is illustrated by point size. Statistical significance of the change ($p < 0.05$) is demarcated by the horizontal dashed line. PTMs: ac = acetyl; me1 = monomethyl; me2 = dimethyl; me3 = trimethyl; prop = propionyl; phos = phosphoryl.
